# Visualization of reactive astrocytes in living brain of Alzheimer’s disease patient

**DOI:** 10.1101/2021.04.13.439744

**Authors:** Min-Ho Nam, Hae Young Ko, Sangwon Lee, Yongmin Mason Park, Seung Jae Hyeon, Woojin Won, Seon Yoo Kim, Han Hee Jo, Jee-In Chung, Young-Eun Han, Gwan-Ho Lee, Yeonha Ju, Thor D. Stein, Mingyu Kong, Hyunbeom Lee, Seung Eun Lee, Soo-Jin Oh, Joong-Hyun Chun, Ki Duk Park, Hoon Ryu, Mijin Yun, C. Justin Lee

**Affiliations:** Center for Neuroscience, Korea Institute of Science and Technology (KIST), Seoul 02792, Republic of Korea; Department of KHU-KIST Convergence Science and Technology, Kyung Hee University, Seoul 02447, Republic of Korea; Department of Nuclear Medicine, Severance Hospital, Yonsei University College of Medicine, Seoul 03722, Republic of Korea; CONNECT-AI Research Center, Yonsei University College of Medicine, Seoul 03722, Republic of Korea; Center for Cognition and Sociality, Institute for Basic Science, Daejeon 34126, Republic of Korea; IBS School, University of Science and Technology, Daejeon 34126, Republic of Korea; KU-KIST Graduate School of Converging Science and Technology, Korea University, Seoul 02841, Republic of Korea; Advanced Analysis Center, KIST, Seoul 02792, Republic of Korea; Boston University Alzheimer’s Disease Research Center and Departments of Neurology and Pathology, Boston University School of Medicine, MA 02130, USA; Molecular Recognition Research Center, KIST, Seoul 02792, Republic of Korea; Virus Facility, Research Animal Resource Center, KIST, Seoul 02792, Republic of Korea; Convergence Research Center for Dementia, KIST, Seoul 02792, Republic of Korea

## Abstract

An early appearance of reactive astrocytes is a hallmark of Alzheimer’s disease (AD)^1,2^, providing a substrate for early diagnostic neuroimaging targets. However, there is no clinically validated neuroimaging probe to visualize the reactive astrogliosis in the human brain *in vivo*. Here, we report that PET/CT imaging with ^11^C-acetate and ^18^F-fluorodeoxyglucose (^18^F-FDG) functionally visualizes the reactive astrocyte-mediated neuronal hypometabolism in the brains with neuroinflammation and AD. We demonstrate that reactive astrocytes excessively absorb acetate through elevated monocarboxylate transporter-1 (MCT1), leading to aberrant GABA synthesis and release which suppresses neuronal glucose uptake through decreased glucose transporter-3 (GLUT3) in both animal and human brains. We propose the non-invasive functional PET/CT imaging for astrocytic acetate-hypermetabolism and neuronal glucose-hypometabolism as an advanced diagnostic strategy for early stages of AD.

## Main

Astrocytes physically and chemically support neighboring neurons under physiological conditions. However, in response to various physical and chemical insults, astrocytes dynamically change their properties morphologically, molecularly, and functionally^3^. The responding astrocytes are termed reactive astrocytes. Reactive astrogliosis, a hallmark of neuroinflammation in Alzheimer’s disease (AD), often precedes neuronal degeneration or death^1,2^ and even directly causes extensive neuronal death^4-6^. Recent studies have also reported that reactive astrocytes aberrantly produce GABA to inhibit neighboring neuronal activity and glucose metabolism^7-10^, which critically contributes to neuronal dysfunction in AD^7,11^. Therefore, *in vivo* imaging of reactive astrogliosis should have a considerable diagnostic value for the early stages of AD. Based on recent reports demonstrating that monoamine oxidase B (MAO-B) is abundantly expressed in the reactive astrocytes of AD^7,10^, the positron emission tomography/computed tomography (PET/CT) of MAO-B has received some endorsement for *in vivo* imaging of reactive astrogliosis^12^. However, several reports have expressed concerns that MAO-B may not be entirely specific for reactive astrogliosis because of its expression in certain types of neurons in the human brain^13-15^. These concerns necessitate the identification of a new molecular target for PET/CT imaging of reactive astrogliosis.

Acetate has been known as an astrocyte-specific energy substrate and an attractive alternative to glucose^16-19^. Acetate has long been believed to be tightly associated with astrocytic metabolism or astrocytic activity^19^, even though there has been a conflicting report^20^. A few previous studies have demonstrated an escalated acetate metabolism in several neuroinflammatory disorders^21,22^. However, whether and how acetate metabolism is associated with astrocyte reactivity has not yet been elucidated. Moreover, acetate has not been clinically considered as PET/CT tracer for AD. In the current study, we investigated the causal relationship between acetate metabolism and astrocyte reactivity and further explored the feasibility of using radioactive acetate as a molecular probe for imaging reactive astrogliosis in AD brain.

The utilization of acetate in astrocytes has been reported to be attributable to a transport through monocarboxylate transporters (MCTs)^17^. Among many subtypes of MCTs, MCT1 is known to be highly expressed in astrocytes^23^ to function as an astrocytic lactate transporter^24^. To examine whether MCT1 functions as a major acetate transporter in astrocytes, we performed the liquid scintillation counting (LSC) of beta-emitting isotopes using ^14^C-acetate with primary cultured astrocytes. We found that treating with SR13800, an MCT1 inhibitor, dose-dependently blocked the ^14^C-acetate uptake in primary cultured astrocytes (Fig. 1a and Extended Data Fig. 1a). The gen-silencing of MCT1 using small-hairpin RNA (shRNA) also significantly blocked the ^14^C-acetate uptake (Fig. 1b and Extended Data Fig. 1b). These findings indicate that MCT1 is essential for acetate uptake in astrocytes.

**Figure 1.**
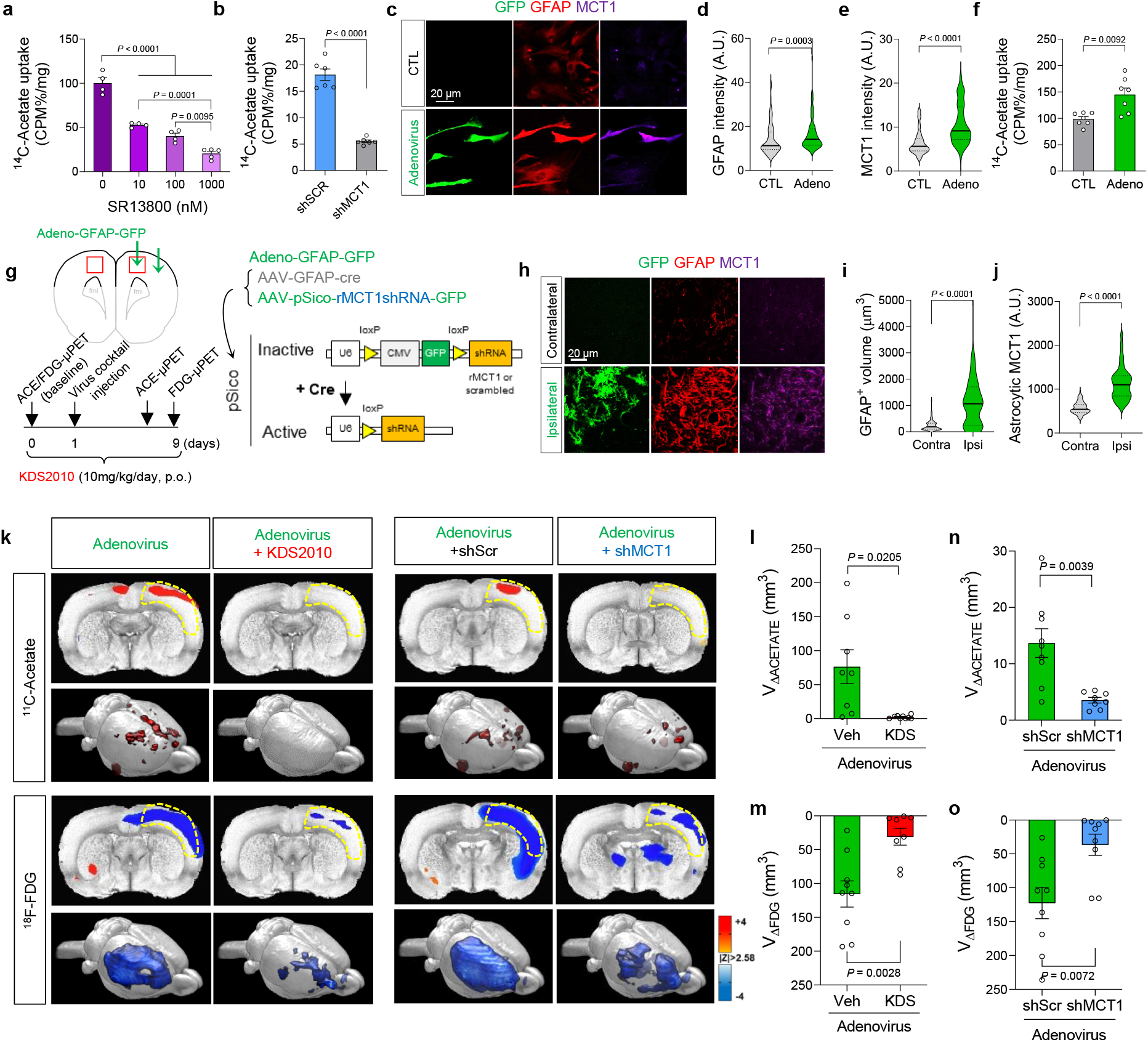
Reactive astrocytes aberrantly uptake acetate through MCT1. **a, b,** Blockade effect of SR13800 (**a**) or *Mct1* gene-silencing (**b**) on ^14^C-acetate uptake in primary cultured astrocytes. **c,** Representative images displaying GFAP and MCT1 expressions in primary cultured astrocytes. **d, e,** Quantification of GFAP and MCT1 immunoreactivity. **f,** The adenovirus effect on ^14^C-acetate uptake. **g,** Schematic diagram of *in vivo* micro-PET imaging of adenovirus model. **h,** Representative images displaying GFAP and MCT1 expressions in adenovirus model. **i, j,** Quantification of the cell volume (**i**) and MCT1 expression (**j**) of GFAP-positive cells. **k,** Left, ^11^C-acetate and ^18^F-FDG PET images in adenovirus model with or without KDS2010 treatment. Right, ^11^C-acetate and ^18^F-FDG PET images in adenovirus model with scrambled-shRNA or MCT1-shRNA. **l, n,** Quantification of the volume of increased ^11^C-acetate uptake. **m, o,** Quantification of the volume of decreased ^18^F-FDG uptake. Mean ± SEM. Significance was assessed by one-way ANOVA with Tukey (**a**), Mann-Whitney test (**d, e, i, j**), or two-tailed unpaired Student’s t-test with (**f, l, n**) or without Welch’s correction (**m, o**).

Next, we investigated whether and how MCT1 expression and acetate uptake are altered in reactive astrocytes. We first validated that adenovirus (Adeno-GFAP-GFP) treatment in primary cultured astrocytes induced reactive astrogliosis, as evidenced by increased expressions of GFAP and MAO-B (Fig. 1c,d and Extended Data Fig. 1c), consistent with previous reports^25^. These reactive astrocytes showed a significantly higher MCT1 expression and increased acetate uptake, compared to control astrocytes (Fig. 1e,f). Two different pro-inflammatory factors using lipopolysaccharide with interferon-γ and amyloid-beta oligomer (Aβ) also increased MCT1 expression in astrocytes, but not in microglia (Extended Data Fig. 1d). The increased MCT1 expression in GFAP-positive reactive astrocytes was also observed in an *in vivo* model of neuroinflammation induced by unilateral adenovirus injection into the motor and somatosensory cortices of the rat brain^8^ (Fig. 1g-j). To investigate whether acetate metabolism is altered in reactive astrocytes and how it impacts neuronal function *in vivo*, we performed microPET scan using ^11^C-acetate followed by ^18^F-FDG with the adenovirus model. ^11^C-acetate as a PET tracer has been widely used in clinics for evaluating myocardial oxidative metabolism and diagnosis of various non-glycolytic tumors^26^. However, ^11^C-acetate has not been tested in the diagnosis of AD. We found a significant increase in ^11^C-acetate uptake and significant decrease in ^18^F-FDG uptake in the ipsilateral cortex of the adenovirus model compared to the contralateral cortex (Fig. 1k-m), consistent with the increased expression of astrocytic MCT1 (Fig. 1h,j and Extended Data Fig. 2) and reduced expression of neuronal glucose transporter 3 (GLUT3) (Extended Data Fig. 3), the main glucose transporter in neurons^27^. These metabolic alterations were prevented by a pharmacological blockade of MAO-B, the key enzyme of reactive astrocytes^7,9,10^, with a selective and reversible MAO-B inhibitor, KDS2010^10^ (Fig. 1k-m). Moreover, the KDS2010 treatment prevented the escalated expression of astrocytic MCT1 in the adenovirus model animals (Extended Data Fig. 2), implying that MCT1 might account for the aberrant acetate uptake. These results indicate that both acetate hypermetabolism and glucose hypometabolism are dependent on MAO-B-mediated neuroinflammation including reactive astrogliosis.

To investigate if astrocytic MCT1 accounts for adenovirus-induced acetate hypermetabolism, we adopted a Cre-LoxP-dependent astrocyte-specific gene-silencing of MCT1 using AAV-GFAP-cre-mCherry and AAV-pSico-rMCT1sh-GFP viruses (Extended Data Fig. 4). We found that the astrocytic gene-silencing of MCT1 significantly reduced the adenovirus-induced ^11^C-acetate uptake, compared to the control scrambled-shRNA (Fig. 1k,n). MCT1 gene-silencing also significantly prevented the adenovirus-induced decrease in ^18^F-FDG uptake (Fig. 1k,o), indicating that MCT1 is necessary for both acetate hypermetabolism and glucose hypometabolism.

We have previously demonstrated that aberrant synthesis and release of GABA from reactive astrocytes causes the neuronal glucose hypometabolism^8^, raising a possibility that the GABA from reactive astrocytes could be responsible for the inverse correlation between acetate and glucose uptake. To test this possibility, we prepared primary cultured cortical astrocytes in the glucose-free astrocyte-media supplemented with both putrescine (180 μM), the key substrate for GABA synthesis in astrocytes, and sodium acetate (1 mM). We then assessed the amount of major metabolites in the putrescine-degradation pathway (Fig. 2a) by performing liquid-chromatography mass-spectroscopy (LC-MS) with the cultured astrocyte homogenate. We found that putrescine treatment significantly increased the level of N-acetyl-GABA and GABA, only when acetate was supplemented (Fig. 2b-d), indicating the critical role of acetate in astrocytic GABA synthesis. Moreover, astrocyte-specific gene-silencing of MCT1 significantly reduced the astrocytic GFAP- and GABA-immunoreactivities in the cortex of the adenovirus model (Extended Data Fig. 5). These results indicate that MCT1-mediated acetate uptake triggers GABA synthesis through the putrescine-degradation pathway in astrocytes. We have also previously demonstrated that Aβ treatment causes reactive astrogliosis and exacerbates astrocytic GABA synthesis^7,10^. To assess the role of acetate in Aβ-induced astrocytic GABA synthesis, we performed the sniffer-patch with GABAc-expressing HEK293T cell as a biosensor (Fig. 2e), as previously described^28^. We found that a 5-day treatment of Aβ (1 μM) turned on the GABA synthesis and Ca^2+^-dependent release, which was significantly increased by the incubation with acetate (10 mM), while acetate alone (without Aβ) did not turn on the GABA synthesis (Fig. 2f,g). These findings indicate that the taken-up acetate boosts Aβ-mediated aberrant GABA synthesis in reactive astrocytes, which could be responsible for neuronal glucose hypometabolism.

**Figure 2.**
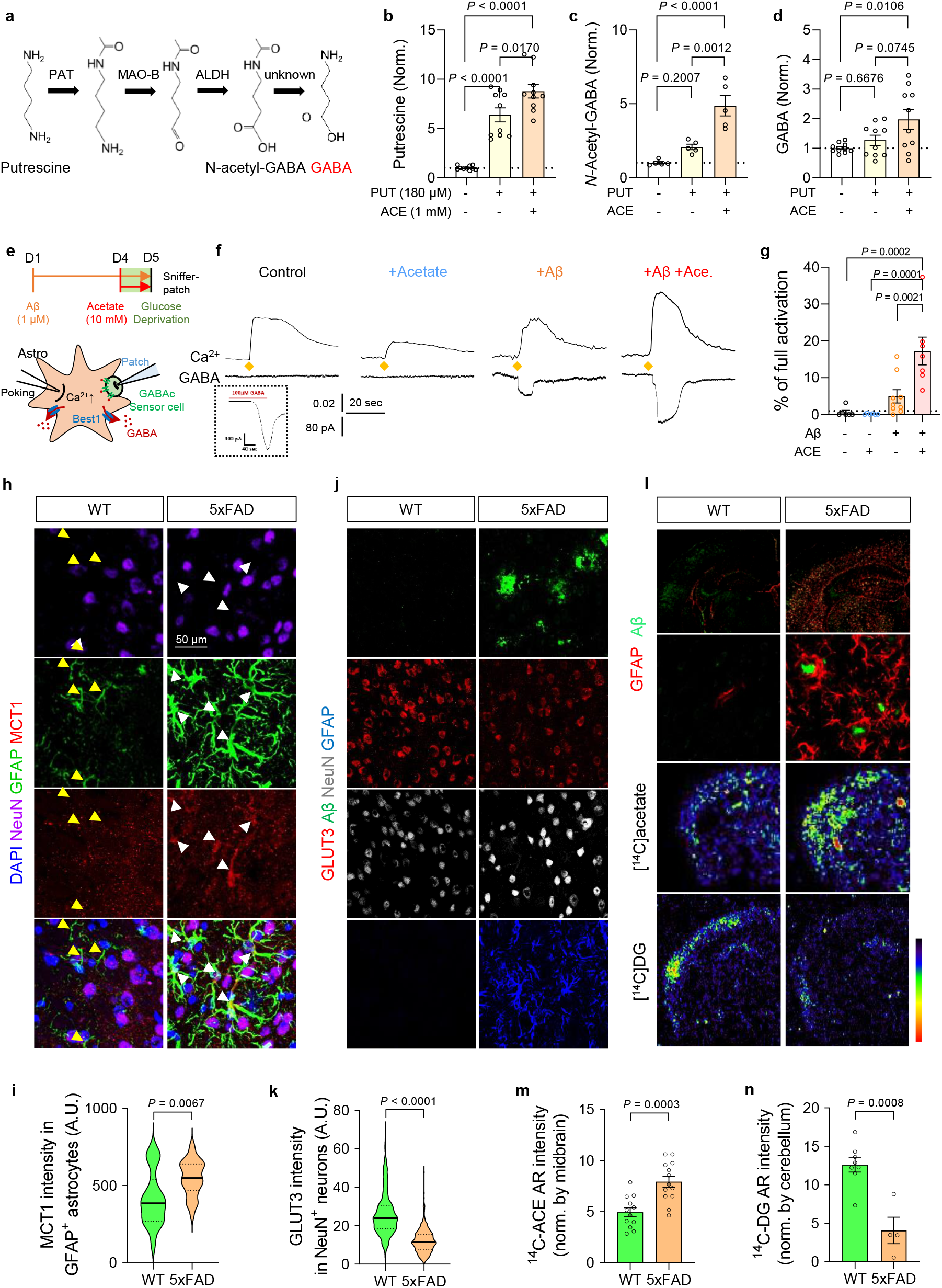
Acetate facilitates astrocytic GABA synthesis to reduce glucose uptake in AD model. **a,** Schematic diagram of astrocytic GABA-synthetic pathway. **b-d**, The level of putrescine, *N*-acetyl-GABA, and GABA analyzed by HPLC. **e,** Schematic diagram of sniffer patch to record GABA current. **f**, Representative traces of Ca^2+^ signal (top) and GABA current (bottom). Yellow diamonds indicate the time-point of poking the astrocyte. **g,** Quantification of poking-induced GABA current. **h,** Representative images displaying GFAP and MCT1 expressions in the cortex of 5xFAD mice. **i,** Quantification of astrocytic MCT1 immunoreactivity. **j,** Representative images displaying NeuN and GLUT3 expressions in the cortex of 5xFAD mice. **k,** Quantification of neuronal GLUT3 immunoreactivity. **l,** Top, representative images displaying Aβ-plaque and GFAP expressions in 5xFAD mice. Bottom, representative autoradiographic images of ^14^C-acetate and ^14^C-DG. **m, n,** Quantification of ^14^C-acetate and ^14^C-DG. Mean ± SEM. Significance was assessed by one-way ANOVA with Tukey (**b, c, d, g**), Mann-Whitney test (**k**) or two-tailed unpaired Student’s t-test (**i, k, m, n**).

We next asked if the similar metabolic alterations can be recapitulated in animal models of AD including 5xFAD and APP/PS1 mice. We found that MCT1 expression in reactive astrocytes was significantly increased in the cortex of both AD models, compared to wild-type littermates (Fig. 2h,i and Extended Data Fig. 6a,b). On the other hand, the neuronal GLUT3 expression was significantly decreased near the amyloid plaques (Fig. 2j,k and Extended Data Fig. 6c,d). Consistently, acetate uptake was significantly increased while glucose uptake was significantly reduced in the cortex of both 5xFAD and APP/PS1 mice, as revealed by autoradiography with ^14^C-acetate and ^14^C-deoxyglucose (^14^C-DG) (Fig. 2l-n and Extended Data Fig. 6e-g). These results indicate that acetate hypermetabolism and glucose hypometabolism are closely associated with increased astrocytic MCT1 and decreased neuronal GLUT3 in animal models of AD, respectively.

To test the clinical relevance, we investigated the protein and mRNA expressions of MCT1 and GLUT3 in the post-mortem hippocampal tissue of AD patients and normal subjects (Extended Data Table 1). Consistent with our previous report^7^, we found abundant reactive astrocytes in the brains with AD, as revealed by higher GFAP immunoreactivity (Extended Data Fig. 7a,b). More importantly, we found that MCT1 was highly localized in GFAP-positive astrocytes, and that the astrocytic MCT1 expression was significantly higher throughout the whole hippocampal formation and frontal cortex of AD patients compared to normal subjects (Fig. 3a-c and Extended Data Fig. 7c,d). MCT1 mRNA level was also significantly higher in AD patients than control subjects (Extended Data Fig. 7e). In contrast, mRNA and protein expressions of GLUT3 in neurons was significantly decreased in AD (Fig. 3h-n and Extended Data Fig. 7g,h), whereas mRNA expression of GLUT1 was not altered in AD (Extended Data Fig. 7i). These results indicate that MCT1 is increased in reactive astrocytes and GLUT3 is reduced in the neighboring neurons, suggesting a feasibility of using ^11^C-acetate and ^18^F-FDG PET/CT in AD patients.

**Figure 3.**
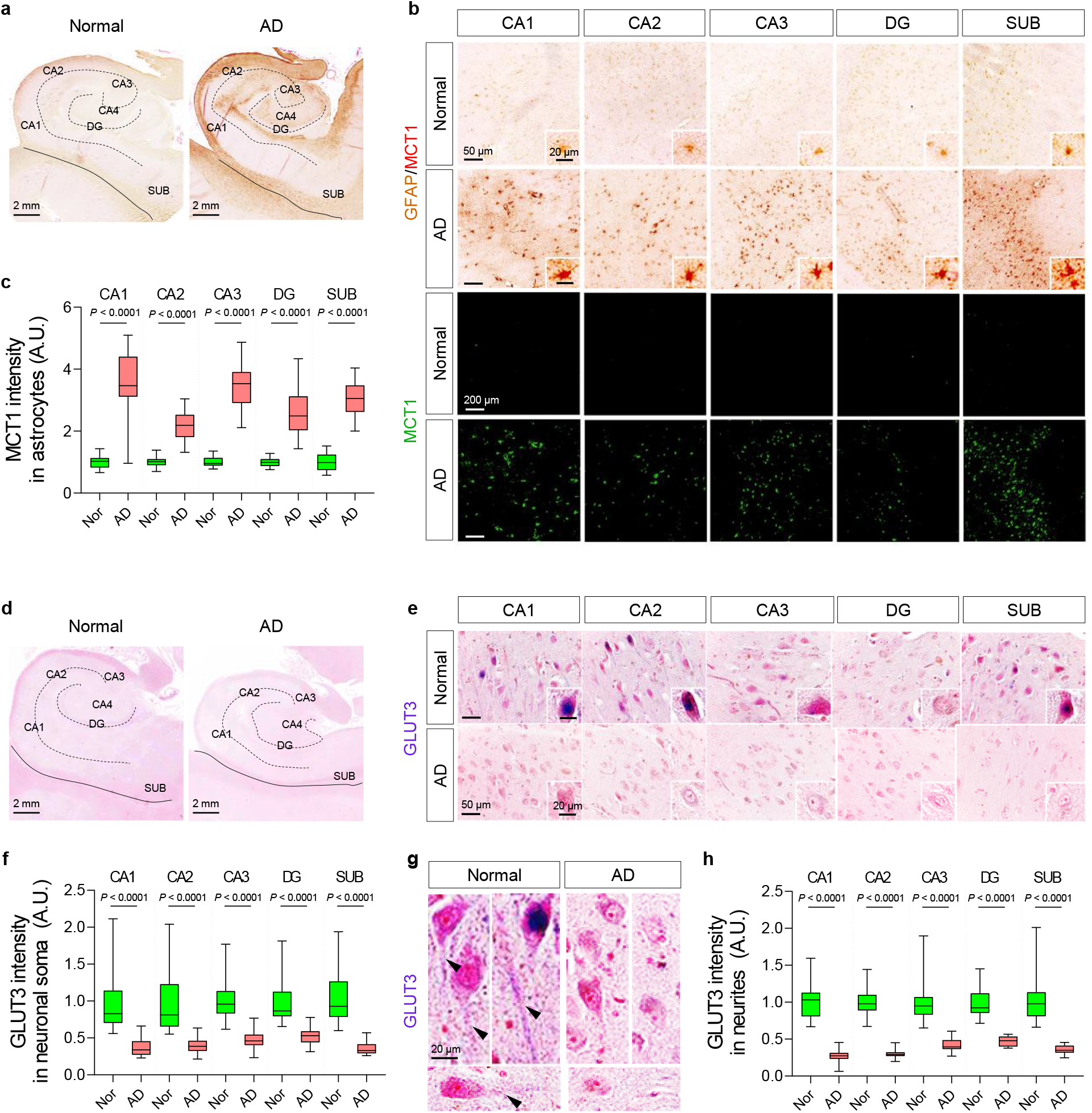
Astrocytic MCT1 is increased while neuronal GLUT3 is decreased in the hippocampus of AD patients. **a, b,** Representative images of double-staining of GFAP and MCT1 in post-mortem hippocampal tissues from normal subjects (N=10) and AD postmortem brains (N=10). **c,** Quantification of astrocytic MCT1 intensity in each hippocampal sub-region. **d, e, g,** Representative images of double-staining of NeuN and GLUT3 in neuronal soma (**d, e**) and neurites (**g**) of post-mortem hippocampal tissues. **f, h,** Quantification of neuronal GLUT3 intensity. Mean ± SEM. Significance was assessed by Mann-Whitney test (**f**, CA1, CA2, DG, and SUB; **h**, SUB) or two-tailed unpaired Student’s t-test with Welch’s correction.

To examine the feasibility, we performed PET/CT imaging with ^11^C-acetate and ^18^F-FDG in 11 AD patients (7 males and 4 females; mean age 77.9 ± 3.6 years) and 10 healthy volunteers (8 males and 2 females; mean age 66.9 ± 9.2 years) (Extended Data Fig. 8). All participants underwent magnetic resonance imaging (MRI) and ^18^F-florbetaben (FBB) PET/CT scans after initial clinical examinations (Extended Data Fig. 8, 9, and Extended Data Table 2). A semi-quantitative analysis of individual PET/CT data was performed in the brain regions implicated in Braak stages of AD, including the entorhinal cortex, hippocampus, fusiform, inferior, middle, and superior temporal gyrus (Fig. 4b). We found that the standardized uptake value (SUV) on ^11^C-acetate PET/CT was significantly higher in all the regions of AD brains (Fig. 4c), while the SUV ratio (SUVR) on ^18^F-FDG PET/CT was significantly decreased (Fig. 4d). To enhance the contrast between the two PET/CT signals, we made subtraction images between ^11^C-acetate and ^18^F-FDG images (Fig. 4a), which allowed better visualization of affected regions. Furthermore, to test if reactive astrogliosis-mediated neuronal hypometabolism is indeed associated with cognitive impairment in AD patients, we analyzed the multiple correlation among ^11^C-acetate uptake, ^18^F-FDG uptake, and cognitive function. We first found that SUV of ^11^C-acetate in each brain region showed significant and strong correlation with either Korean-Mini Mental State Examination (K-MMSE) score (Fig. 4e and Extended Data Fig. 10) or memory score in Seoul Neuropsychological Screening Battery (SNSB) (Extended Data Fig. 11). Similar high correlations were observed in the SUVR of ^18^F-FDG (Extended Data Fig. 10). Finally, multiple linear regression showed MMSE scores were highly correlated with both ^11^C-acetate SUV and ^18^F-FDG SUVR with the highest R^2^ values in the entorhinal cortex and hippocampus (Fig. 4f). Taken together, these results indicate that the reactive astrogliosis visualized by ^11^C-acetate and the associated neuronal dysfunction visualized by ^18^F-FDG are highly correlated with cognitive impairment of AD patients. Furthermore, our findings provide the first *in vivo* evidence for the critical role of reactive astrogliosis in human AD symptomatology, which has been suspected for several decades based on animal studies.

**Figure 4.**
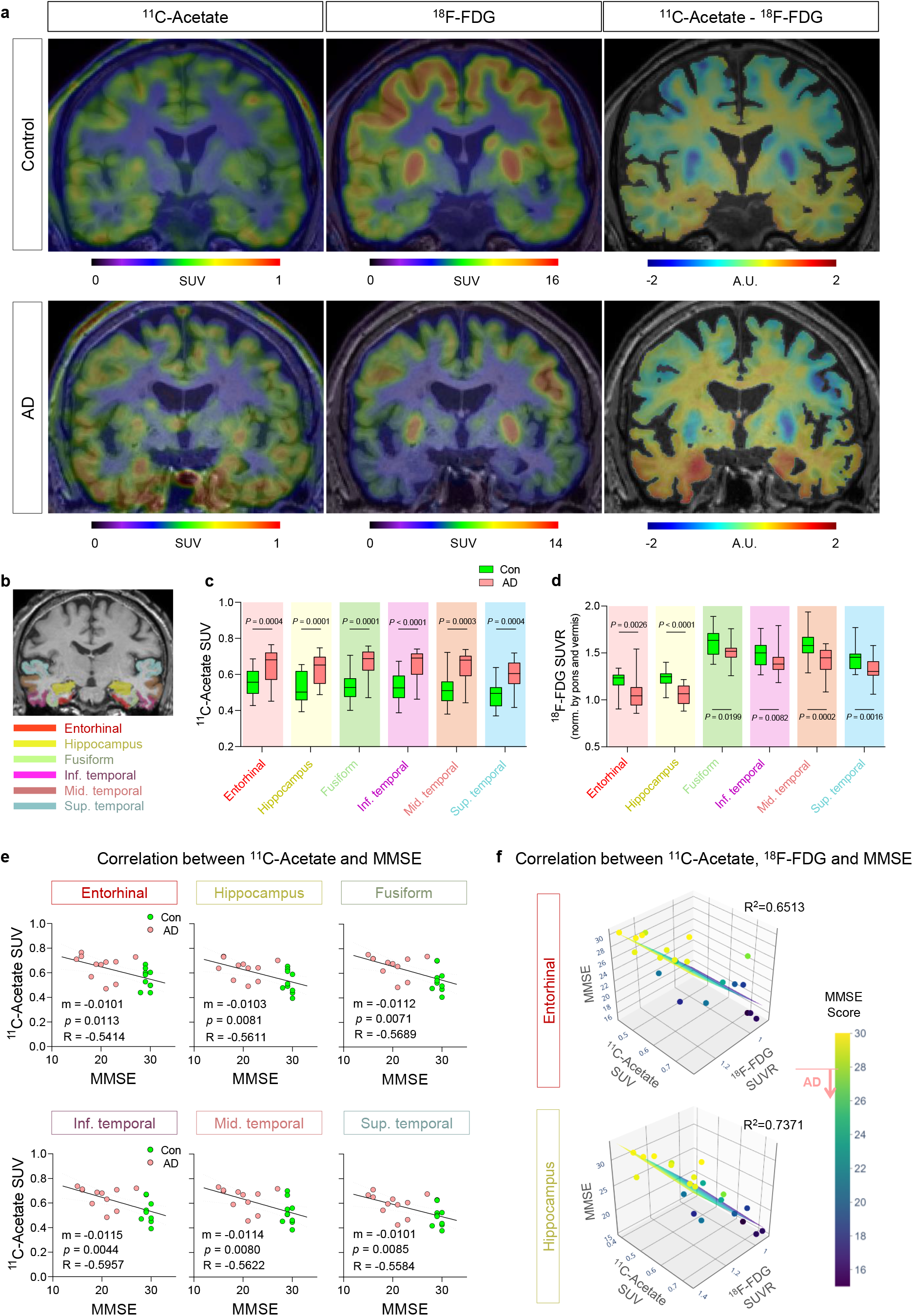
^11^C-acetate and ^18^F-FDG PET/CT imaging for visualizing reactive astrogliosis in AD patients’ brains. **a,** Representative PET/CT images of ^11^C-acetate and ^18^F-FDG and their subtraction images in control and AD patients. **b,** ROIs from an MR image. **c**, **d,** Quantification of ^11^C-acetate and ^18^F-FDG SUV in each ROI of control (N=10) and AD patients (N=11). **e,** Correlation between ^11^C-acetate SUV and MMSE scores. **f,** Multiple correlations between ^11^C-acetate SUV, ^18^F-FDG SUVR, and MMSE scores. Mean ± SEM. Significance was assessed by two-tailed unpaired Student’s t-test (**c, d**) or linear regression (**e**) or multiple linear regression (**f**) with Pearson’s correlation.

In this study, we have delineated the molecular and cellular mechanisms of acetate hypermetabolism and glucose hypometabolism in AD by identifying the molecular targets of astrocytic MCT1 for acetate uptake and neuronal GLUT3 for glucose uptake (Extended Data Fig. 12). We show the feasibility of using ^11^C-acetate and ^18^F-FDG PET/CT for *in vivo* imaging of reactive astrocytes and altered neuronal metabolism in AD. The proposed tools and concepts should be further optimized and utilized to develop advanced diagnostics for reactive astrogliosis in early stages of AD.

## Acknowledgments

This study was supported by IBS-R001-D2 from the Institute for Basic Science funded by the Korean Ministry of Science and ICT to C.J.L.; NRF-2018M3C7A1056898 from National Research Foundation (NRF) of Korea to M.Y.; NRF-2018M3C7A1056894, NRF-2020M3E5D9079742, and KIST Grants (2E30320, 2E30762) to H.R.; and NRF-2018M3C7A1056897 to M.-H.N. The schematic figure (Extended Data Fig. 12) was drawn using Biorender.

## Author contributions

Conceptualization by M.-H.N., M.Y., and C.J.L.; Methodology by S.E.L. and G.H.L.; Investigation and validation by M.-H.N, H.Y.K., S.L., Y.M.P., S.J.H., W.W., S.Y.K., H.H.J., J.I.C., Y.H., Y..J., M.K., and H.L.; Resources by M.-H.N, T.D.S., H.L., S.J.O., K.D.P., J.-H.C, H.R., M.Y., C.J.L; Writing - Original Draft by M.-H.N.; Writing - Review & Editing by M.-H.N., H.Y.K., H.R., M.Y., and C.J.L.; Supervision by H.R., M.Y., and C.J.L.; Funding acquisition by M.-H.N., H.R., M.Y., and C.J.L.

## Competing interests

The authors declare that they have no conflict of interest.

## Data and materials availability

All data is available in the main text or the supplementary materials upon any reasonable requests.

## Notes

### Competing Interest Statement

The authors have declared no competing interest.

