## Supplementary material for "Visualization of reactive astrocytes in living brain of Alzheimer’s disease patient": methods, Extended Fig. 1 to 12, Table 1, 2

**Online methods**

Animals

Adult male Sprague Dawley rats (Koatech, Korea) weighing 250–300 grams (8 to 10-week-old) were used for experiments with adenovirus-induced neuroinflammation models. Adult male APP/PS1 (11 to 12-month-old) and 5xFAD transgenic mice (6 to 8-month-old) were used for experiments for investigating the metabolic change of reactive astrocytes. The animal experimental procedures were approved by Institutional Animal Care and Use Committee of Korea Institute of Science and Technology (KIST; Seoul, Korea; approval No. KIST-2019-042), and Yonsei University (Seoul, Korea; approval No. 2017-0187). The animals were kept on a 12 hr light-dark cycle with controlled temperature (21 ± 1°C) and humidity (50 ± 10%) and had ad libitum access to food and water. Animal care was followed by National Institutes of Health (NIH) guidelines.

Drug administration

KDS2010, a MAO-B inhibitor, was synthesized as previously described (Park et al., 2019). KDS2010 was administered by dissolving the compound in drinking water for 9 days after baseline micro-PET imaging. The amount of KDS2010 was calculated as 10 mg/kg daily.

Virus preparation and injection

To induce neuroinflammation including reactive astrocytosis, we used the Adeno-GFAP-GFP virus which is developed previously [1]. To induce adenovirus-induced neuroinflammation, we injected Adeno-GFAP-GFP into two points in the sensory cortex (AP = -2.0 mm, ML = +2.0 mm and +4.0 mm, DV = -1.5 mm and -2.5 mm from bregma) using stereotaxic apparatus under under general anesthesia with 2% isoflurane. For gene-silencing of MCT1, we prepared MCT1-shRNA whose sequences of complementary oligomers were 5′-TGC TCC ACT TAA TCA GGC TTT CTT CAA GAG AGA AAG CCT GAT TAA GTG GAG CTT TTT TC-3′ and 5′-TCG AGA AAA AAG CTC CAC TTA ATC AGG CTT TCT CTC TTG AAG AAA GCC TGA TTA AGT GGA GCA-3′. The knockdown efficiency was tested by reverse transcription polymerase chain reaction (RT-PCR) with cDNA from rat primary cultured astrocytes which were electroporated with the shRNA vector. For AAV-based shRNA expression, a lentiviral vector containing the MCT1-shRNA gene was constructed into the HpaI–XhoI restriction enzyme sites of the pSico AAV vector in an AAV-DJ capsid. AAV-DJ was engineered via DNA family shuffling technology which created a hybrid capsid from eight AAV serotypes. AAV-DJ displays a higher transduction efficiency in vitro than any wild-type serotype. AAV-DJ is known to successfully infect a broad range of various cell types *in vivo*[2]. The viral vectors were purified by iodixanol gradients by the KIST Virus Facility. The minimum number of viral particles was $1.0 \times 10^{12}$ genome copies (GC)/mL, which is a concentrated virus package.

To selectively knockdown astrocytic MCT1, we injected AAV-GFAP-Cre-mCherry and $AAV_{DJ}$ containing pSico-MCT1-shRNA-GFP (or pSico-scrambled-shRNA-GFP for control) into the same points in the sensory cortex using stereotaxic apparatus. A mixture of 1 μL of Adenovirus for inducing neuroinflammation, 0.5 μL of AAV-GFAP-Cre-mCherry, and 0.5 μL of pSico-MCT1-shRNA-GFP for knockdown astrocytic MCT1, was slowly injected at the target sites with a rate of 0.15 ml/min using 33G Hamilton syringe connected to an UltraMicroPump (WPI). After injection, the needle was left in place for an additional 7 min before being slowly retracted.

Primary Astrocyte Culture

The cerebral cortex of a P1 pup was dissected free of adherent meninges, minced, and dissociated into single-cell suspension by trituration through a Pasteur pipette. Dissociated cells were plated onto either 12-mm glass coverslips or six-well plates coated with 0.1 mg/mL poly d-lysine (PDL; Sigma). Cells were grown in Dulbecco's modified Eagle's medium (DMEM; Gibco) supplemented with 25 mM glucose, 10% heat-inactivated horse serum, 10% heat-inactivated fetal bovine serum, 2 mM glutamine, and 1,000 units/mL penicillin-streptomycin. 3 days later, cells were vigorously washed with repeated pipetting using medium and the media was replaced to get rid of debris and other floating cell types.

In vitro $^{14}$C-acetate uptake assay in primary astrocyte

To measure $^{14}$C-acetate uptake, primary astrocytes were seeded in 24-well plates and incubated with a standard culture medium for 24 hours. For the inhibition test of SR13800 (Tocris), the cells were incubated in 0, 10, 100 and 1000 nM of SR13800 with culture medium at 37°C for 12 hours. Subsequently $^{14}$C-Acetate (PerkinElmer, USA) 0.5 μCi/0.5 mL of external solution (150 mM NaCl, 3 mM KCl, 10 mM HEPES, 22 mM Sucrose, 2 mM $MgCl_2$, 2 mM $CaCl_2$, pH 7.4) were added to each wells and the cells were incubated at 37°C for 20 minutes. At the end of the uptake period, each well was washed twice with ice-cold PBS. The cells were lysed with 200 μL of 0.2 N NaOH for 2 h at room temperature. After the addition of a scintillation cocktail (Ultima Gold; PerkinElmer), the radioactivity was measured by a liquid scintillation counter (Tri-Carb, PerkinElmer, USA). The measured radioactivity was normalized to protein concentration which was performed using a BCA Protein Assay kit (Thermo Fisher Scientific). Experiments were performed three times using three replicates for each experimental condition.

Western blotting

The cells were lysed with 1% sodium dodecyl sulfate (SDS) lysis buffer containing protease inhibitor cocktail (Roche, Germany), and total protein concentration was determined by protein assay (Thermo Fisher Scientific). Equal amounts (5 μg) of protein from each sample were separated by SDS-polyacrylamide gel electrophoresis (SDS-PAGE, Bio-rad) and transferred to polyvinylidene difluoride membranes (Millipore, USA). The membranes were blocked with 5% skim milk at room temperature for 1 h and then incubated with rabbit anti-MAOB (1:1000, Novus), rabbit anti-MCT1 (1:1000, Alomone), and mouse anti-beta actin (1:2000, Invitrogen) at 4°C overnight. Membranes were washed in TBS-T and incubated with goat anti-rabbit or anti-mouse IgG horseradish peroxidase (1:2000, GeneTex) as the secondary antibody. The antigen-antibody complexes were visualized using the ECL western blotting substrate (Thermo Fisher Scientific).

microPET imaging

Each rat was scanned a total of four times: baseline $^{11}C$-acetate and baseline $^{18}F$-FDG scans prior to adenovirus injection, $^{11}C$-acetate scan, and $^{18}F$-FDG scan at 8 days after adenovirus injection. KDS2010 (10mg/kg/day) was administered for 9 days after the baseline PET image. Rat imaging was performed using a microPET scanner (Inveon, Siemens Healthcare, Germany), which has a transaxial resolution of 1.4 mm full width at half maximum and a 12.7 mm field of view. Animals were anesthetized with 2.5% isoflurane before the administration of radiotracer. A dose of 37 MBq (1 mCi) of $^{11}C$-acetate was injected into the tail vein of the rats. Following the 20 min of uptake time, $^{11}C$-acetate PET data were acquired for 40 min under 2% isoflurane anesthesia. For the $^{18}F$-FDG PET scan, the rats were administered with 20 MBq (0.54 mCi) of $^{18}F$-FDG and allowed to uptake for 40 min on a heating pad under 2% isoflurane anesthesia. Then, 40 min static acquisition was performed for $^{18}F$-FDG PET scan. All PET data were reconstructed with three-dimensional (3D) ordered subset expectation-maximization (OSEM) with 2 iterations and 18 subsets. The voxel size was 0.776 × 0.776 × 0.796 mm and the matrix size was 128 × 128 × 159.

MicroPET image processing was performed using the Analysis of Functional NeuroImages (AFNI) software[3]. $^{11}C$-acetate and $^{18}F$-FDG PET images were registered to the MRI template of Sprague Dawley rat brain using an automated template-based registration algorithm and manual manipulation for minor misalignment[4]. To statistically compare $^{11}C$-acetate and $^{18}F$-FDG PET images in the adenovirus-induced neuroinflammation models (with and without KDS2019 treatment, and with and without scrambled-shRNA and MCT1-shRNA), a voxel-wise t-test was performed using 3dttest in AFNI software. The 3-D rendering images fused T1-weighted MR image with the statistical parametric map were displayed using MRIcroGL software (https://www.mccauslandcenter.sc.edu/mricrogl/).

Human brain samples

Neuropathological examination of normal subject and AD human brain samples was determined using procedures previously established by the Boston University Alzheimer's Disease Center (BUADC). Next of kin provided informed consent for participation and brain donation. Institutional review board approval for ethical permission was obtained through the BUADC center. This study was reviewed by the Institutional Review Board of the Boston University School of Medicine (Protocol H-28974) and was approved for exemption because it only included tissues collected from post-mortem subjects not classified as human subjects. The study was performed in accordance with institutional regulatory guidelines and principles of human subject protection in the Declaration of Helsinki. The sample information is listed in Supplementary Table S1.

Immunostaining for confocal microscopy

For immunohistochemistry, sections were first incubated for 1.5 h in a blocking solution (0.3% Triton-X, 2% goat serum, and 2% donkey serum in 0.1 M PBS) and then immunostained with a mixture of primary antibodies in a blocking solution at 4°C. Primary antibodies used are as follow: chicken anti-GFAP (1:500, ab5541, Millipore), rabbit anti-MCT1 (1:200, ab3538p, Millipore) for rat samples, rabbit anti-MCT1 (1:200, AMT-011, Alomone) for mouse samples, mouse anti-NeuN (1:1000, MAB377, Millipore), mouse anti-GLUT3 (1:200, sc-74399, Santa Cruz) for rat samples, rabbit anti-GLUT3 (1:200,

AGT-023, Alomone) for mouse samples, and guinea pig anti-GABA (1:300, ab175, Millipore). After extensive washing, sections were incubated with corresponding fluorescent secondary antibodies for 2 h and then washed with PBS 3 times. If needed, DAPI (1:3,000, Pierce) staining was performed. To visualize amyloid plaques, sections were incubated in 1 mM thioflavin-S, which had been dissolved in 50% ethanol for 8 min. Sections were rinsed with 80% ethanol twice for differentiation and washed with PBS 3 times. Finally, sections were mounted with a fluorescent mounting medium (S3023, Dako) and dried. A series of fluorescent images were obtained with an A1 Nikon confocal microscope, and Z-stack images in 3-μm steps were processed for further analysis using or NIS-Elements (Nikon, Japan) software and ImageJ program (NIH, MD, USA). Any alterations in brightness or contrast were equally applied to the entire image set. Specificity of primary antibody and immunoreaction was confirmed by omitting primary antibodies or changing fluorescent probes of the secondary antibodies.
For immunocytochemistry, we fixed the cultured astrocytes with 4% paraformaldehyde at 4°C for 10 min. After washing with 0.1 M PBS three times, we performed immunocytochemistry according to the same procedures as immunohistochemistry.

Double staining immunohistochemistry for the human postmortem brain

First staining: Paraffin-embedded tissues were sectioned in a coronal plane at 10 to 20 mm. Endogenous alkaline phosphatase was blocked using 3% hydrogen peroxide in TBS. Sections were blocked with 2.5% normal horse serum (Vector Laboratories) for 1 hr and then incubated with MCT1- or GLUT3-specific antibody for 24 hr. After washing, sections were incubated with ImmPRESS-AP anti-rabbit IgG (alkaline phosphatase) polymer detection reagent (Vector Laboratories: MP-5402) for 30 minutes at room temperature. MCT1 signals were developed with a Vector Red alkaline phosphatase substrate kit (Vector Laboratories). GLUT3 signals were developed with a Vector Blue substrate kit (Vector Laboratories: SK-5300).

Second staining: To verify the localization of MCT1 in reactive astrocytes, mouse monoclonal antibody to GFAP (1:200 dilution; Santa Cruz Biotechnology, Dallas, TX, USA) was incubated over the MCT1-stained tissue slides for 24 hr. After washing three times with PBS, the slides were processed with Vector ABC Kit (Vector Laboratories, Inc., Burlingame, CA, USA). The GFAP immunoreactive signals were developed with DAB chromogen (Thermo Fisher Scientific, Meridian, Rockford, IL, USA). Otherwise, GLUT3-stained slides were subsequently counterstained with Vector Nuclear Fast Red (Vector Laboratories: H-3403). Double-stained tissue slides were processed back to xylene through an increasing ethanol gradient [70%, 80%, and 95% (1 time), and 100% (2 times)] and then mounted.

Sniffer patch

The sensor cells were originated from HEK293T cells with stable GABAc sensor cell line construction. The construction is described as followed. The full length of GABAc receptor (GABAc) CDS was cloned into pHR-CMV-IRES-EmGFP lentiviral expression vector (a gift from A. Radu Aricescu, Addgene plasmid # 113888) via EcoRI and AgeI restriction sites. For lentiviral particle production, the cloned plasmids were transiently transfected into HEK293T cells with packaging plasmid psPAX2 and envelope plasmid pMD2.G using Lipofectamine 3000 transfection reagent (Thermo Fisher, USA) according to the manufacturer's instructions. At 24 h and 48 h post-transfection, the cell culture media containing lentivirus particles were harvested, filtered through 0.45 μm-pore-size filters, and stored at -80°C until needed. For the generation of GABAc-expressing stable cell lines, lentiviral particles-containing media were directly overlaid on HEK293T cells in the presence of polybrene (Merck, Germany) at a final concentration of 4 μg/ml. After 24 h, the supernatant was changed with fresh medium and cultured for 2 days. GFP-positive cells were single-cell sorted into 96-well plates using a MoFlo Astrios Cell Sorter (Beckman Coulter, USA) and cultured another 2~3 weeks to allow for clonal expansion. HEK293T cells were regularly tested for mycoplasma contamination.

The day before the sniffer patch, cortical astrocytes were seeded from culture dishes onto 12 mm glass coverslips coated with PDL in 24-well plates. On the day of the sniffer patch, sensor cells were seeded from culture dishes onto astrocyte-placed coverslips. Astrocytes, which were co-cultured with GABAc receptor sensor cell, were incubated with 5 mM Fura-2AM (mixed with 5 μL of 20% Pluronic acid; P3000MP, Invitrogen) for 40 min and washed at room temperature and subsequently transferred to a microscope stage for imaging. External solution contained (in mM): 150 NaCl, 10 HEPES, 3 KCl, 2 $CaCl_2$, 2 $MgCl_2$, pH adjusted to pH 7.3 and osmolality to 320-325 mOsm $kg^{-1}$. For $Ca^{2+}$ imaging, Intensity images of 510 nm wavelength were taken at 340 nm and 380 nm excitation wavelengths using CoolLED (pE-$340^{fura}$). Astrocytic $Ca^{2+}$ responses were induced by poking as previously described [5]. Two resulting images were used for ratio calculations in Axon Imaging Workbench version 9.0 (Axon Instruments). GABAcR-mediated currents from sensor cell were recorded under voltage clamp (Vh = -50 mV) using Multiclamp 700B amplifier (Molecular Devices), acquired with pClamp 11.0.3 Recording electrodes (4-7 MΩ) were filled with (mM): 140 CsCl, 0.5 $CaCl_2$, 10 HEPES, and 10 EGTA (pH adjusted to 7.3 with CsOH and Osmolality with 285–295 mOsm $kg^{-1}$). To normalize the different expressions of GABAc in sensor cells, 100 μM of GABA bath application was performed to obtain maximal GABAc current from each sensor cells. Sniffed current, which is mediated by released GABA from astrocytes, was divided by maximal GABAc current.

<u>Autoradiography</u>

APP/PS1 and 5xFAD mice (4 mice per each group) were intravenously injected with $^{14}$C-acetate (3 μCi) or $^{14}$C-DG (3 μCi) in 200 μL of saline and perfused with 4% PFA at 1hour postinjection of tracers, respectively. The brains were quickly removed and frozen. For autoradiography, coronal sections (20-μm thickness) were prepared using a cryostat at −20°C and mounted on poly-L-lysine–coated slides. The sections were exposed to an imaging plate (BAS-IP SR2025, Fuji Film, Japan) for 2 weeks. The plates were visualized using a bio-imaging analyzer system (Typhoon FLA 7000, GE Healthcare, USA). The intensity of radioactivity in the ipsilateral cortex was quantified using imageJ (NIH). The intensity was normalized with the intensity in the midbrain and cerebellum for $^{14}$C-acetate and $^{14}$C-DG, respectively.

<u>LC-MS</u>

Drug-treated cortical astrocytes (DIV 12-13) were detached from culture dishes by incubating 2.5% trypsin (15140-122, Gibco). Trypsinized astrocytes were washed with Dulbecco's phosphate-buffered saline (DPBS, LB001-02, Welgene). Washed astrocyte pellets were perfectly dried by suction and stored at -80℃.

Metabolites were extracted from the rat primary cultured astrocytes by adding 100 μL cold methanol:water (7:3), containing GABA $d_2$ (final concentration of 0.5ppm) as an internal standard. The samples were lysed by performing three cycles of freeze-thaw using liquid nitrogen, and the samples were vortexed for 30 sec and then centrifuged at 20,817 × *g* (14,000 rpm) for 10 min. The supernatant was dried with nitrogen gas using Turbovap LV (Biotage). The samples were then reconstituted with the mobile phase consisting of 80% A and 20% B. The samples were analyzed using a UPLC-MS/MS instrument consisting of an ExionLC AD system (AB Sciex) and a triple-quadrupole 4500 mass spectrometer (AB Sciex) equipped with an electrospray ionization source. An Acquity UPLC® BEH HILIC column (2.1 mm x 100 mm, 1.7 μm, Waters) was used to separate the metabolites with the mobile phase A (0.1% formic acid in acetonitrile) and B (0.1% formic acid and 50 mM ammonium formate in water). The flow rate was set to 0.4 mL/min and the injection volume was 5 μL with the following gradient program: starting at 20% B, was maintained for 7 min, and a linear gradient was initiated to reach 80% B over 30 sec, then maintained for 1 min, then decreased to 20% B over 30 sec and maintained at 20% for 1 min. The total run time was 10 min.

The mass spectrometer was operated in positive ion multiple reaction monitoring (MRM) mode with the following parameters: curtain gas (CUR) at a pressure of 30 psi, turbo IonSpray voltage (IS) at 5500 V, source temperature (TEM) at 550°C, declustering potentials (DP) at 26 V, entrance potentials (EP) at 10 V, collision energies (CE) at 15 V, collision cell exit potential (CXP) at 8 V. Data acquisition was processed using the Analyst® Software (AB Sciex).

Human PET/CT imaging

Eleven patients were clinically diagnosed with AD who presented at the dementia outpatient clinic of Severance Hospital, Yonsei University Health System. AD was diagnosed according to the criteria of the National Institute of Neurological and Communicative Disorders and Stroke and the Alzheimer's Disease and Related Disorders Association (NINCDS-ADRDA)[6] and the guideline proposed by Petersen and colleagues[7]. Ten subjects, who had no previous history of neurological disorders or subjective symptoms of cognitive impairment, were selected for normal control (CON). All subjects underwent neuropsychological testing (Seoul Neuropsychological Screening Battery, SNSB), brain MRI, and $^{18}$F-FDG, $^{18}$F-florbetaben (FBB), and $^{11}$C-Acetate PET/CT. This study was approved by the institutional review board of Severance Hospital (IRB No. 4-2018-1070), and written informed consent was obtained from all participants. All patients fasted at least 6 hours before $^{11}$C-acetate and $^{18}$F-FDG PET/CT scans, which were performed using Discovery 600 (General Electric Healthcare, Milwaukee, MI, USA). A dose of 740 MBq (20 mCi) of $^{11}$C-acetate was intravenously administered to the patients. For the $^{18}$F-FDG PET/CT scan, 4.1 MBq (0.11 mCi) per body weight (kg) of $^{18}$F-FDG was intravenously administered to the patients. After the 10-min of uptake period for $^{11}$C-acetate and 40-min for $^{18}$F-FDG, PET/CT scans were performed. The scan time of $^{11}$C-acetate and $^{18}$F-FDG were 20-min and 15-min, respectively. The information of CT scan for attenuation correction was following: 0.5 seconds rotation time, 200 mA, 120 kVp, 3.75 mm section thickness, 10.0 mm collimation, and 9.375 mm table feed per rotation. The acquired PET data were reconstructed using the OSEM with 2 iterations and 32 subsets.

Image processing was performed using MATLAB (The Mathworks, Inc, Natick, MA, USA)-based software called Statistical Parametric Mapping (SPM12, Wellcome Trust Centre for Neuroimaging, London, UK). $^{18}$F-FDG PET/CT and corresponding T1-weighted images were co-registered to the $^{11}$C-acetate PET/CT using an automatic registration algorithm based on mutual information. The co-registered T1 images were segmented into gray, white matter and cerebral spinal fluid (CSF) using SPM12's segmentation algorithm. $^{11}$C-acetate and the co-registered $^{18}$F-FDG PET/CT images were normalized for mean counts in the brain within each scan. The co-registered and normalized $^{18}$F-FDG PET/CT images were subtracted from the normalized $^{11}$C-acetate PET/CT to generate a difference image between the two data sets. A semi-quantitative analysis was performed using the FreeSurfer pipeline 6.0 (Massachusetts General Hospital, Harvard Medical School; http://surfer.nmr.mgh.harvard.edu/). T1-weighted MR images were used to segment cortical region-of-interest (ROI) based on the Desikan-Killiany Atlas. For both $^{11}$C-acetate and $^{18}$F-FDG PET/CT images, the SUV images were calculated as follows; (decay-corrected activity [kBq] per tissue volume [mL])/(injected FDG activity [kBq] per body mass [g]). The $^{18}$F-FDG PET/CT images were intensity normalized by the mean value of the pons and vermis to derive a standardized uptake value ratio (SUVR).

Quantitative real-time RT-PCR

Total RNAs were isolated from the frozen brain tissues using TRIzol reagent (MRC, TR118). Fifty nanograms of RNA was used as a template for quantitative RT-PCR amplification, using SYBR Green Real-time PCR Master Mix (Toyobo, QPK-201, Osaka, Japan). Primers were standardized in the linear range of the cycle before the onset of the plateau. Human GAPDH was used as an endogenous control to standardize the amount of RNA in each reaction. The following sequences of primers were used. *MCT1* forward: 5´-TAC CTC CAG ACT CTC CTG GC -3´; *MCT1* reverse: 5´- GTC CCC TCC GCA AAG TCT AC -3´; *GAPDH* forward: 5'- GAA ATC CCA TCA CCA TCT TCC-3' and reverse: 5'-GAG GCT GTT GTC ATA CTT CTC-3'.

Image quantification

Confocal microscopic images were analyzed using the ImageJ program (NIH) and Imaris 9 (Bitplane). For measurement of GFAP and MCT1 immunoreactivity in astrocytes, we first threshold the binary GFAP+ image to define single astrocytes as region-of-interests (ROIs) using ImageJ. Then we measured the intensity of GFAP or MCT1 in every ROI from 8-bit GFAP+ or MCT1+ images. For measurement of GFAP+ volume, we made the surface for each GFAP+ cell with GFAP+ images using Imaris, and then we collected the mean intensity values of the volume of each ROI. GFAP, MCT1, and GLUT3 immunoreactivities in human hippocampal tissues were also analyzed using the ImageJ program (NIH).

Statistical Analysis

Statistical analyses were performed using Prism 9 (GraphPad Software, Inc.). Differences between two different groups were analyzed with the two-tailed Student's unpaired t-test. For assessment of change of a group by a certain intervention, the significance of data was assessed by the two-tailed Student's paired t-test. For comparison of multiple groups, one-way analysis of variance (ANOVA) with Tukey's or Dunnett's multiple comparison test, or two-way ANOVA with Bonferroni's multiple comparison test was assessed. For assessing the correlations between two factors, linear regression was performed and Pearson's correlation coefficient was calculated. For assessing the correlations between three factors, multiple linear regression was performed and the MMSE score or SNSB (memory) score was chosen as the outcome. The normality of the distribution of each dataset was tested. When the data does not normally distributed we performed appropriate non-parametric tests such as Mann-Whitney test or Kruskal-Wallis ANOVA test. For comparisons of two or multiple groups, we also tested if the variances are statistically different across the groups. If the variance is different, appropriate corrections were applied to the statistical tests. $P < 0.05$ was considered to indicate statistical significance throughout the study. The significance level is represented as asterisks (*$P < 0.05$, **$P < 0.01$, ***$P < 0.001$; ns, not significant). Unless otherwise specified, all data are presented as mean ± SEM. No statistical method was used to predetermine sample size. Sample sizes were determined empirically based on our previous experiences or the review of similar experiments in literatures. The numbers of animals used are described in the corresponding figure legends or on each graph. All experiments were done with at least three biological replicates. Experimental groups were balanced in terms of animal age, sex and weight. Animals were genotyped before experiments, and they were all caged together and treated in the same way. Prior to administration of virus injection or drug administration, animals were randomly and evenly allocated to each experimental group. The data analysis of animal experiments was performed by two independent investigators. However, investigators were not blinded to outcome assessments.

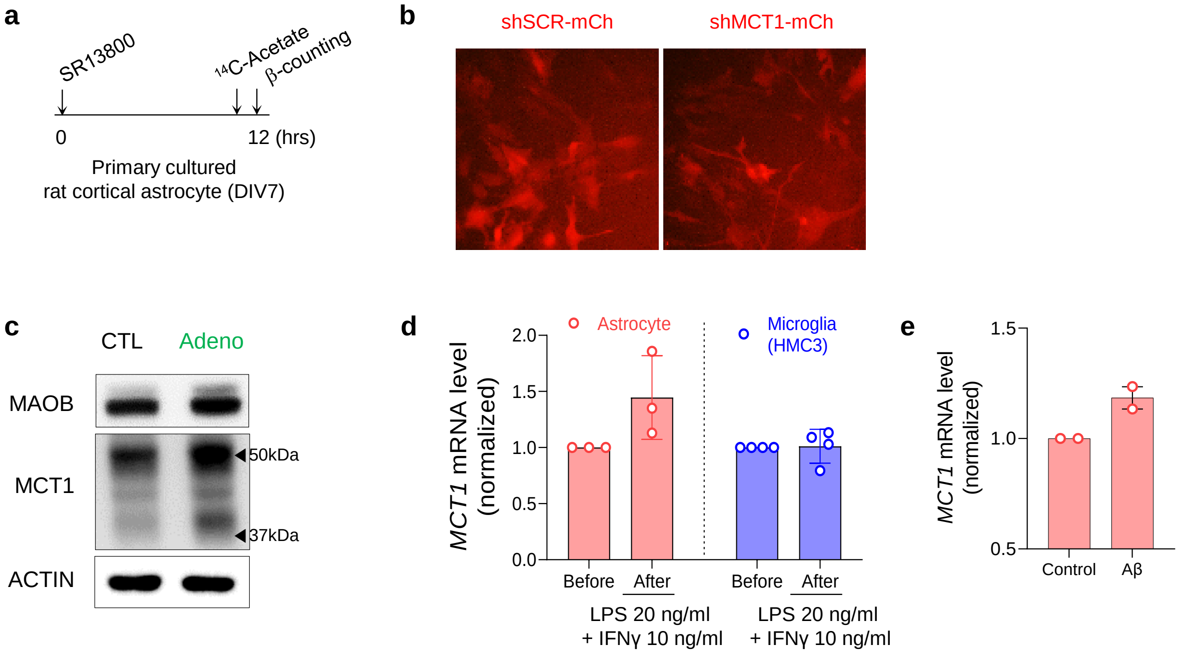


**Extended Data Figure 1. The augmented expression level and function of MCT1 in reactive astrocytes**
**a,** Timeline of LSC experiments with primary cultured astrocytes. **b,** Representative fluorescent images demonstrating the expressions of shRNA-mCherry. **c,** Western blotting of MAOB and MCT1 in adenovirus-treated reactive astrocytes. **d,** MCT1 mRNA expression in LPS and IFNγ-induced reactive astrocytes and reactive microglia. **e,** MCT1 mRNA expression in control and Aβ-treated astrocytes

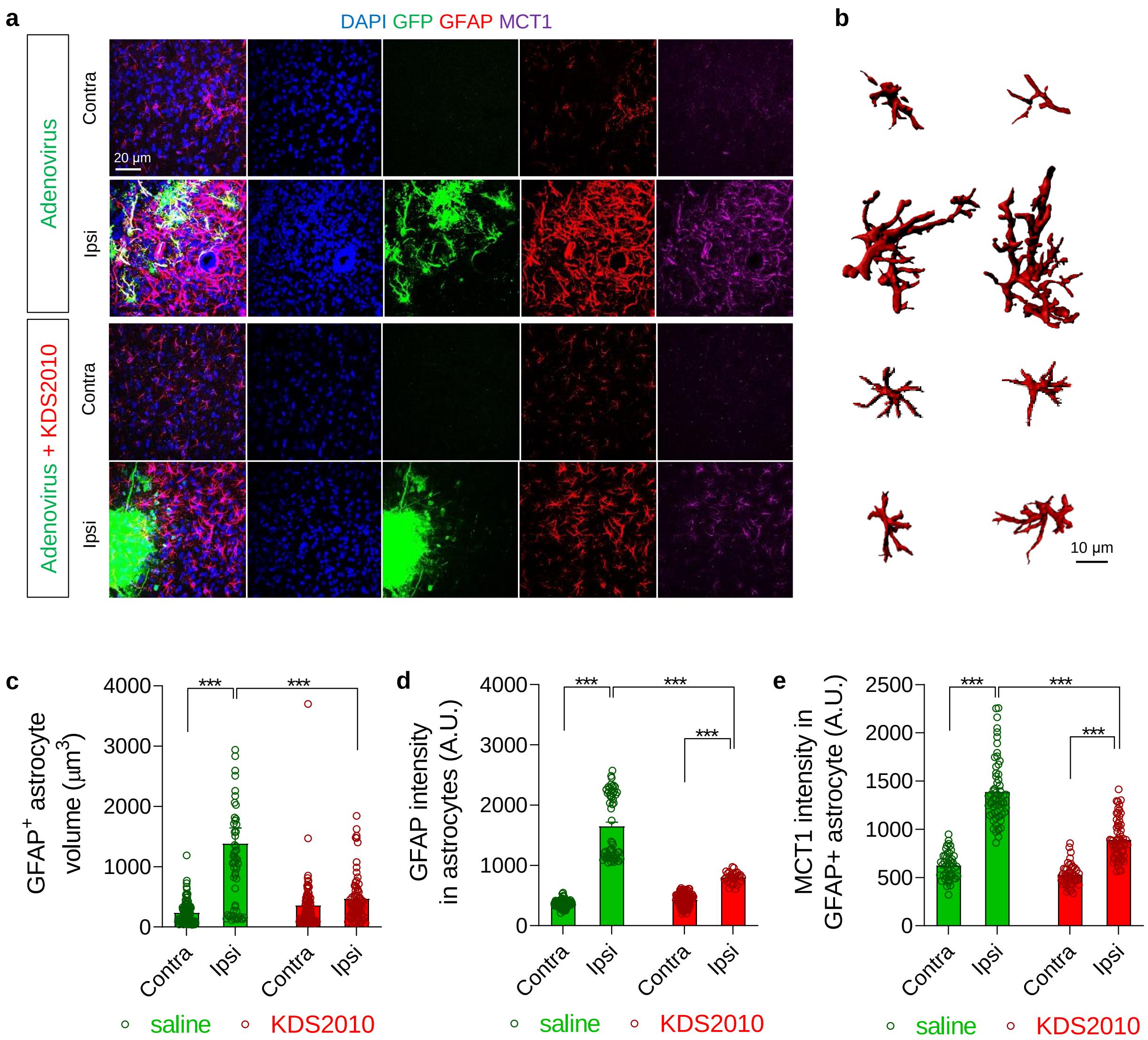


**Extended Data Figure 2. Adenovirus increased astrocyte reactivity and MCT1 expression level, which were attenuated by KDS2010 treatment.**
**a,** Representative confocal images of rat cortical tissues. **b,** Representative 3D-rendered GFAP-positive astrocytes. **c,** Quantification of the volume of GFAP-positive astrocytes. Adenovirus increased the volume of GFAP-positive astrocytes, which was significantly attenuated by KDS2010 treatment. **d,** Quantification of the GFAP intensity in astrocytes. Adenovirus increased the GFAP intensity, which was significantly attenuated by KDS2010 treatment. **e,** Quantification of the MCT1 immunoreactivity in astrocytes. Adenovirus increased the MCT1 immunoreactivity, which was significantly attenuated by KDS2010 treatment.

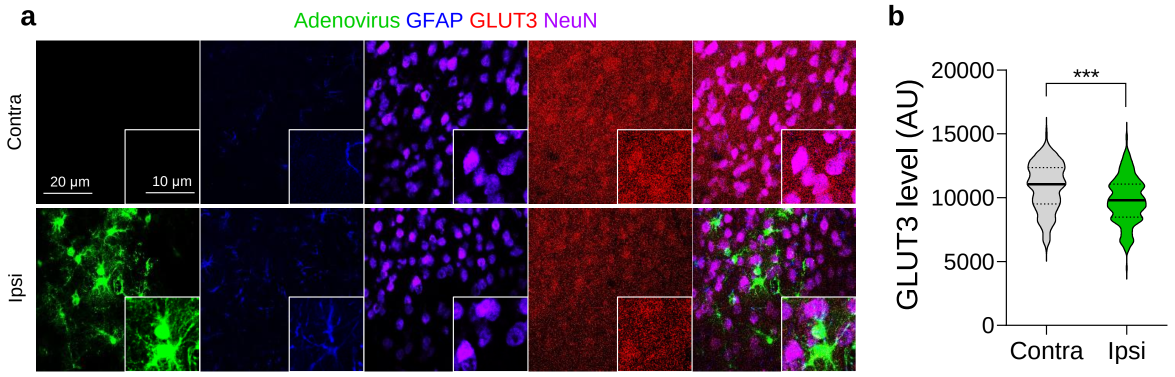


**Extended Data Figure 3. GLUT3 expression is reduced by adenovirus injection.** (**A**) Representative confocal images of rat cortical tissues demonstrating reduced GLUT3 expressions by adenovirus infection. **b,** Quantification of GLUT3 expression.

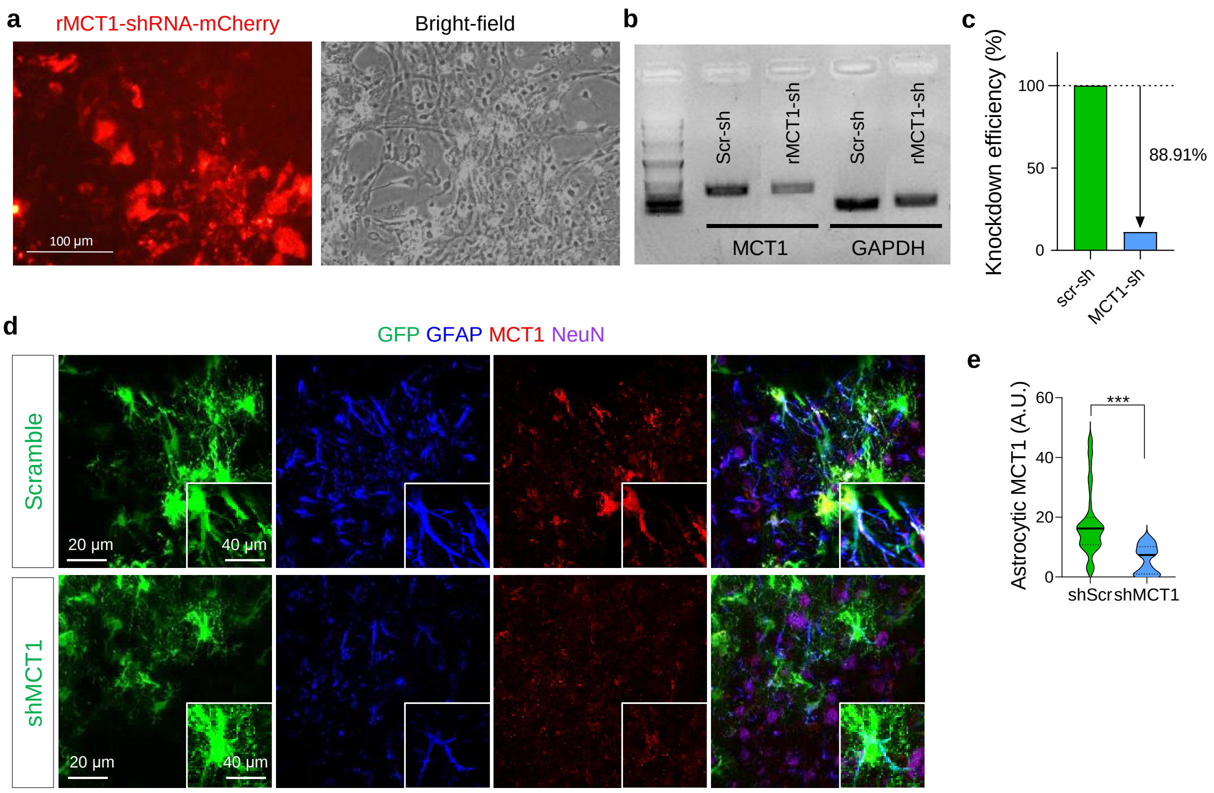


**Extended Data Figure 4. In vitro and in vivo knockdown efficiency of MCT1-shRNA.**

**a,** Representative images of primary cultured rat cortical astrocytes. **b,** RT-PCR bands demonstrating MCT1-shRNA markedly reduced MCT1 expression. **c,** Quantification of knockdown efficiency of MCT1 from qRT-PCR experiments. **d,** Representative confocal images of rat cortical tissues demonstrating reduced MCT1 expressions in the MCT1-shRNA-expressed astrocytes. **e,** Quantification of MCT1 expression in astrocytes.

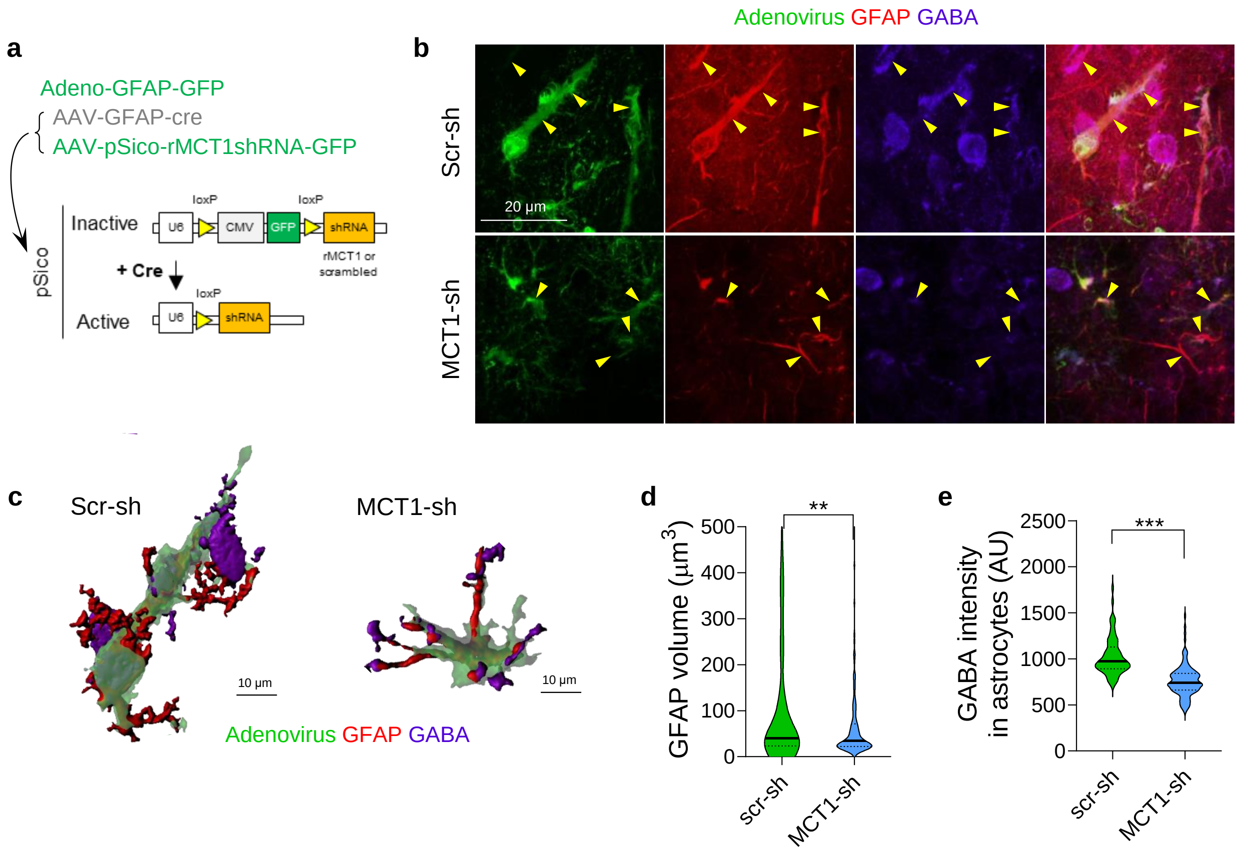


**Extended Data Figure 5. Astrocyte-specific MCT1 gene silencing ameliorates reactive astrogliosis in adenovirus model.**
**a,** Schematic diagram of cre-dependent astrocyte-specific MCT1 gene-silencing. **b,** Representative confocal images of rat cortical tissues demonstrating reduced GFAP and GABA expressions in the MCT1-shRNA-expressed astrocytes. **c,** Representative 3D-rendered astrocytes-expressing adeno-GFAP-GFP virus, GFAP, and GABA. **d,** Quantification of GFAP-positive volume of each astrocyte. **e,** Quantification of GABA expression in each astrocyte.

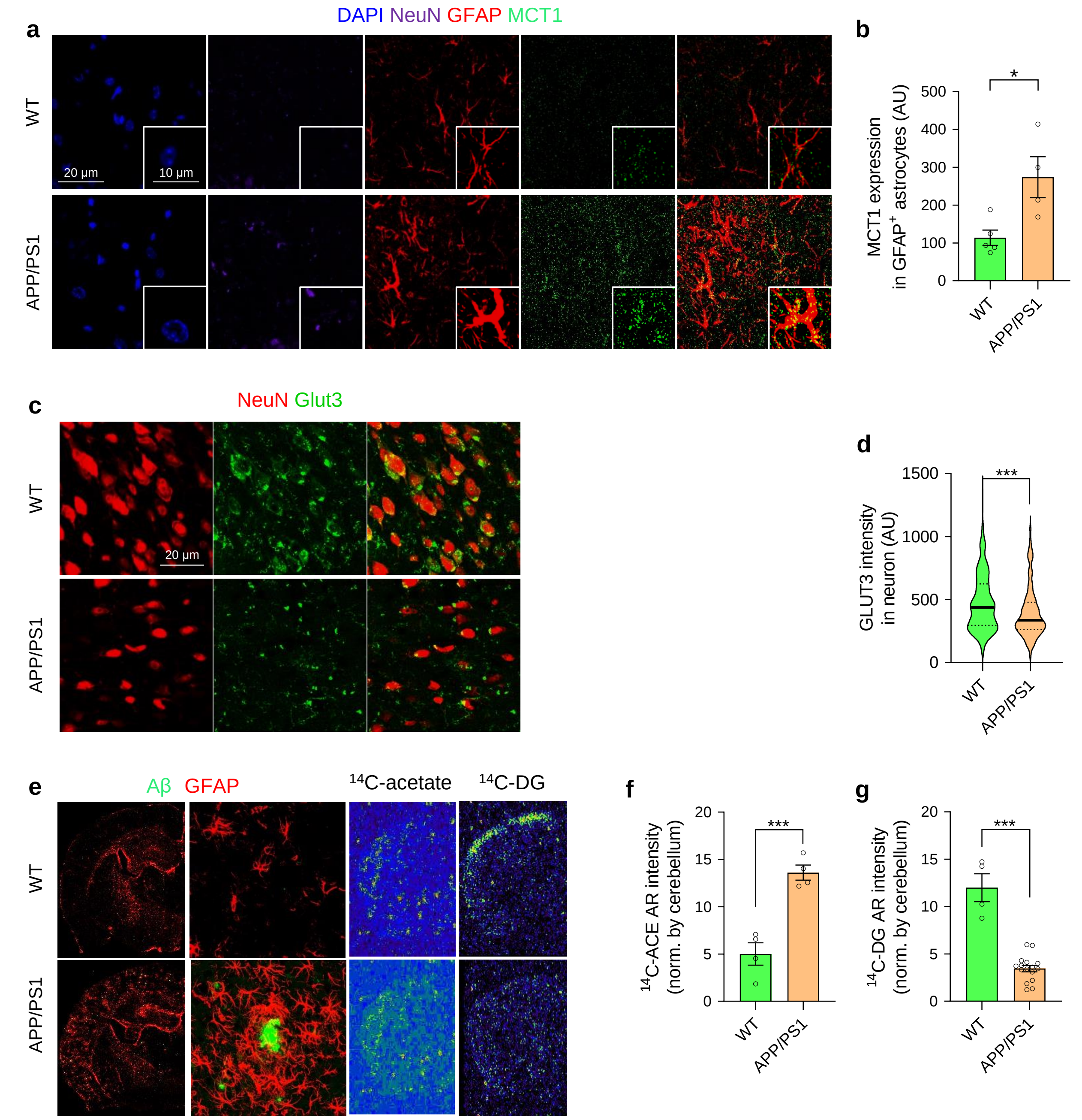


**Extended Data Figure 6. MCT1 level and acetate uptake is increased, while GLUT3 level and glucose uptake is decreased in APP/PS1 transgenic mice.**

**a,** Representative confocal images of APP/PS1 mouse cortical tissues demonstrating increased astrocytic MCT1 expressions. **b,** Quantification of GFAP-positive astrocytic MCT1 expression. **c,** Representative confocal images of APP/PS1 mouse cortical tissues demonstrating decreased neuronal GLUT3 expressions. **d,** Quantification of neuronal GLUT3 expression. **e,** Left, representative confocal images of Aβ and GFAP in APP/PS1 transgenic mice and the littermates. Right, representative autoradiography images of $^{14}$C-acetate and $^{14}$C-DG. **f,** Quantification of $^{14}$C-acetate intensity from autoradiography images. **g,** Quantification of $^{14}$C-DG intensity from autoradiography images.

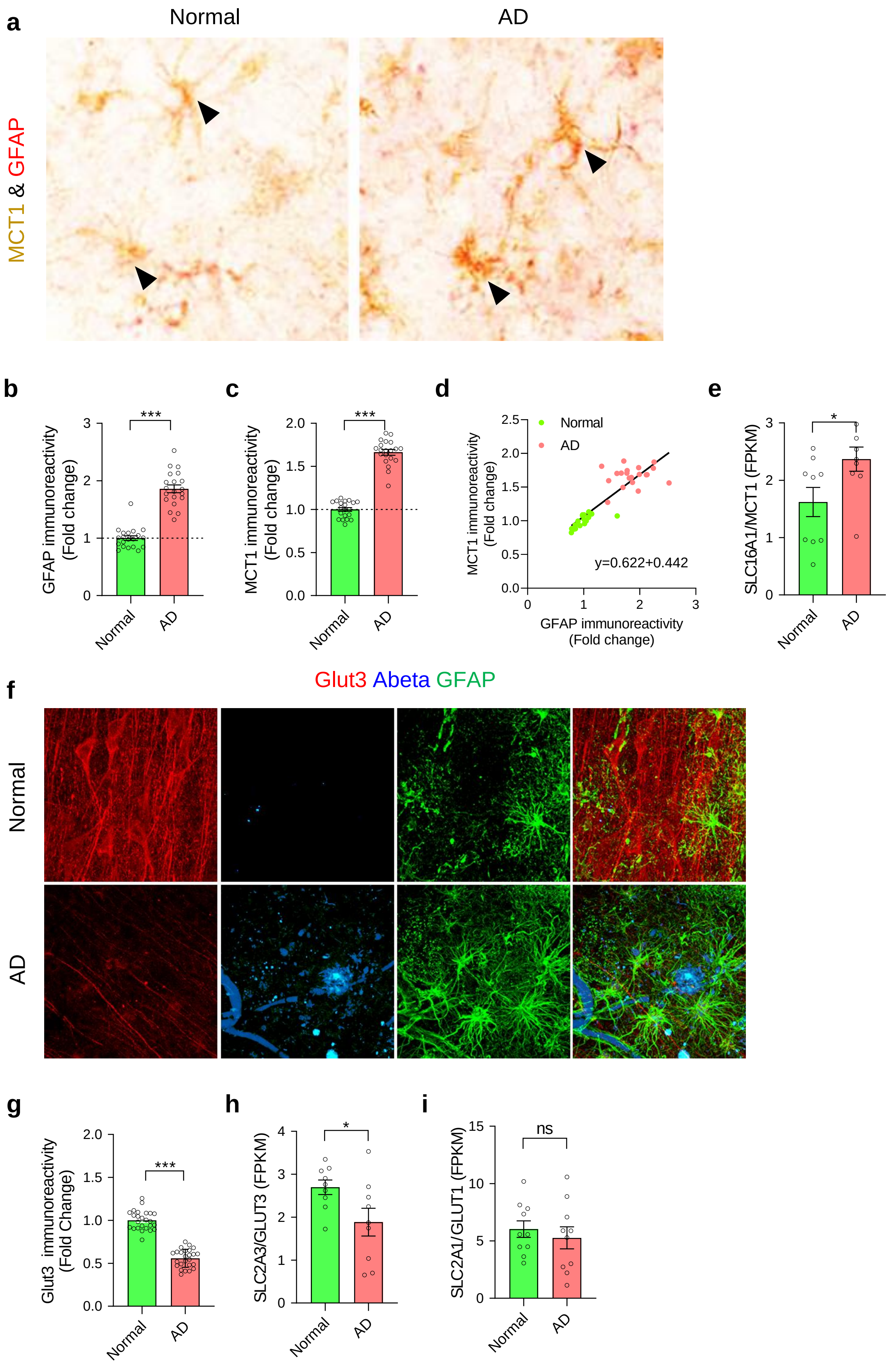

a
Normal
AD
MCT1 & GFAP
b
GFAP immunoreactivity (Fold change)
***
Normal
AD
c
MCT1 immunoreactivity (Fold change)
***
d
Normal
AD
MCT1 immunoreactivity (Fold change)
GFAP immunoreactivity (Fold change)
y=0.622+0.442
e
SLC16A1/MCT1 (FPKM)
*
f
Glut3 Abeta GFAP
Normal
AD
g
Glut3 immunoreactivity (Fold Change)
***
h
SLC2A3/GLUT3 (FPKM)
*
i
SLC2A1/GLUT1 (FPKM)
ns

**Extended Data Figure 7. Increased MCT1 and decreased GLUT3 levels in the forebrain of post-mortem brain tissues from AD patients and normal subjects.** **a,** Representative images of human post-mortem forebrain tissues double-immunostained with MCT1 and GFAP. **b,** Quantification of GFAP immunoreactivity. **c,** Quantification of MCT1 immunoreactivity. **d,** A positive correlation between MCT1 and GFAP immunoreactivities. **e,** *SLC16A1* mRNA (encoding MCT1) expression in the frontal cortex of normal subjects and AD patients. **f,** Representative images of human post-mortem forebrain tissues immunostained with GLUT3 and GFAP. **g,** Quantification of GLUT3 immunoreactivity. (**H**) *SLC2A3* mRNA (encoding GLUT3) expression in the frontal cortex of normal subjects and AD patients.

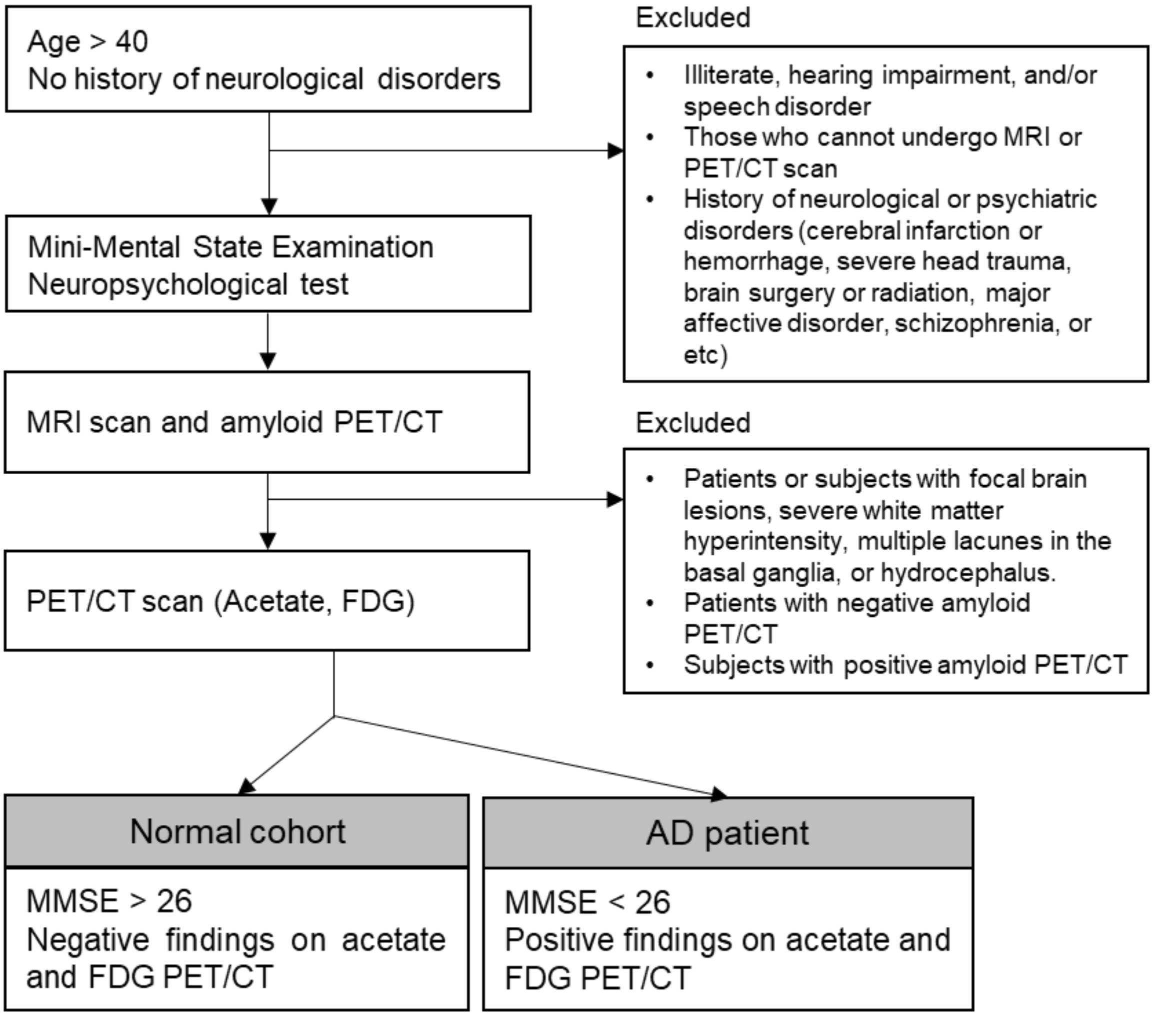


**Extended Data Figure 8. Human subject selection criteria and procedure**

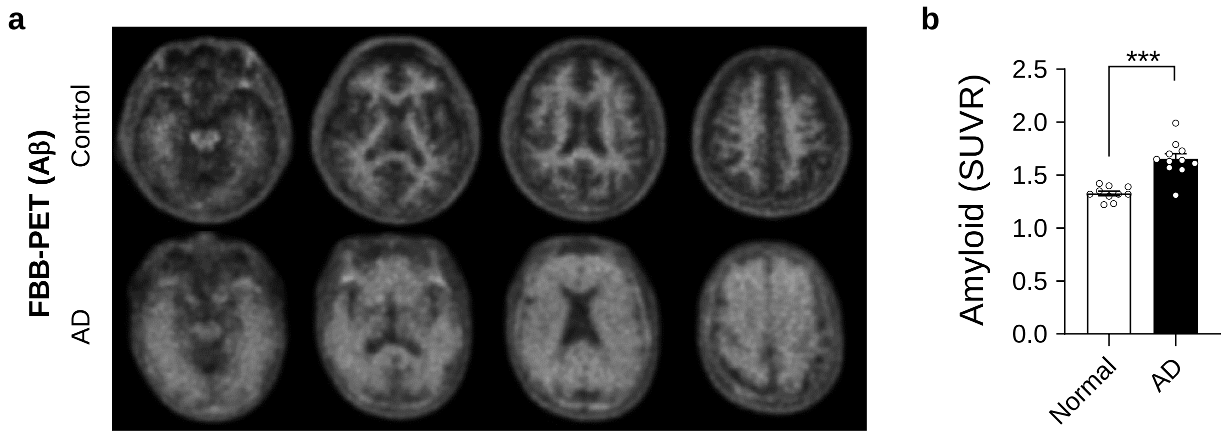


**Extended Data Figure 9. Diagnostic screening by FBB-PET.**
**a,** Representative brain FBB-PET images of Aβ in control subjects and AD patients **b,** SUVR of Aβ in control subjects and AD patients.

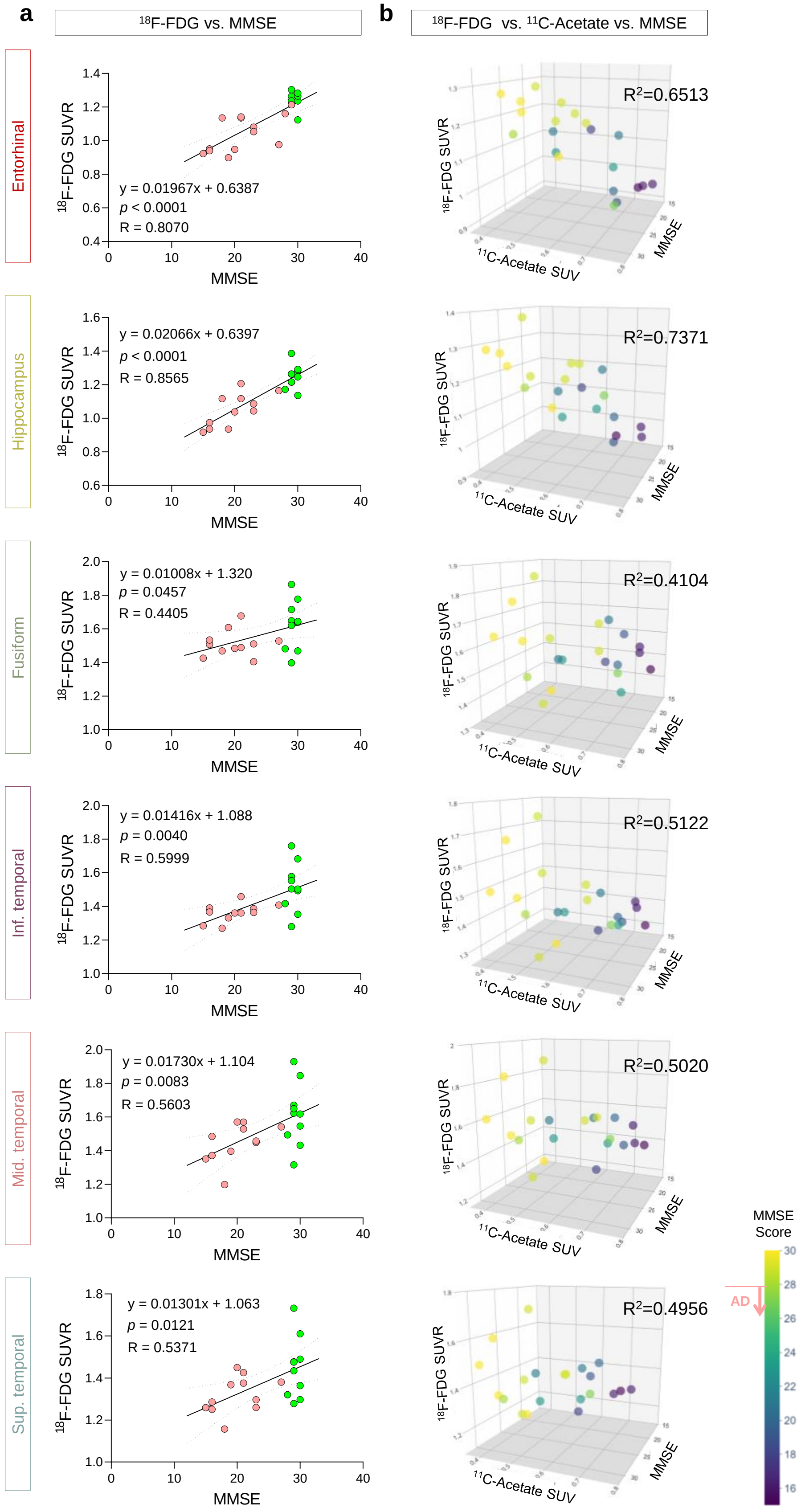
a
18F-FDG vs. MMSE
b
18F-FDG vs. 11C-Acetate vs. MMSE
Entorhinal
y = 0.01967x + 0.6387
p < 0.0001
R = 0.8070
R2=0.6513
Hippocampus
y = 0.02066x + 0.6397
p < 0.0001
R = 0.8565
R2=0.7371
Fusiform
y = 0.01008x + 1.320
p = 0.0457
R = 0.4405
R2=0.4104
Inf. temporal
y = 0.01416x + 1.088
p = 0.0040
R = 0.5999
R2=0.5122
Mid. temporal
y = 0.01730x + 1.104
p = 0.0083
R = 0.5603
R2=0.5020
Sup. temporal
y = 0.01301x + 1.063
p = 0.0121
R = 0.5371
R2=0.4956
18F-FDG SUVR
MMSE
11C-Acetate SUV
MMSE Score
AD

**Extended Data Figure 10. Correlations between $^{18}$F-FDG SUVR, $^{11}$C-acetate SUV, and MMSE score in various human brain region.**

**a,** In every region-of-interest, a significant correlation between $^{18}$F-FDG SUVR MMSE score was observed. **b,** The multiple linear regression analysis showed that $^{18}$F-FDG SUVR and $^{11}$C-acetate SUV has marked correlations with MMSE scores.

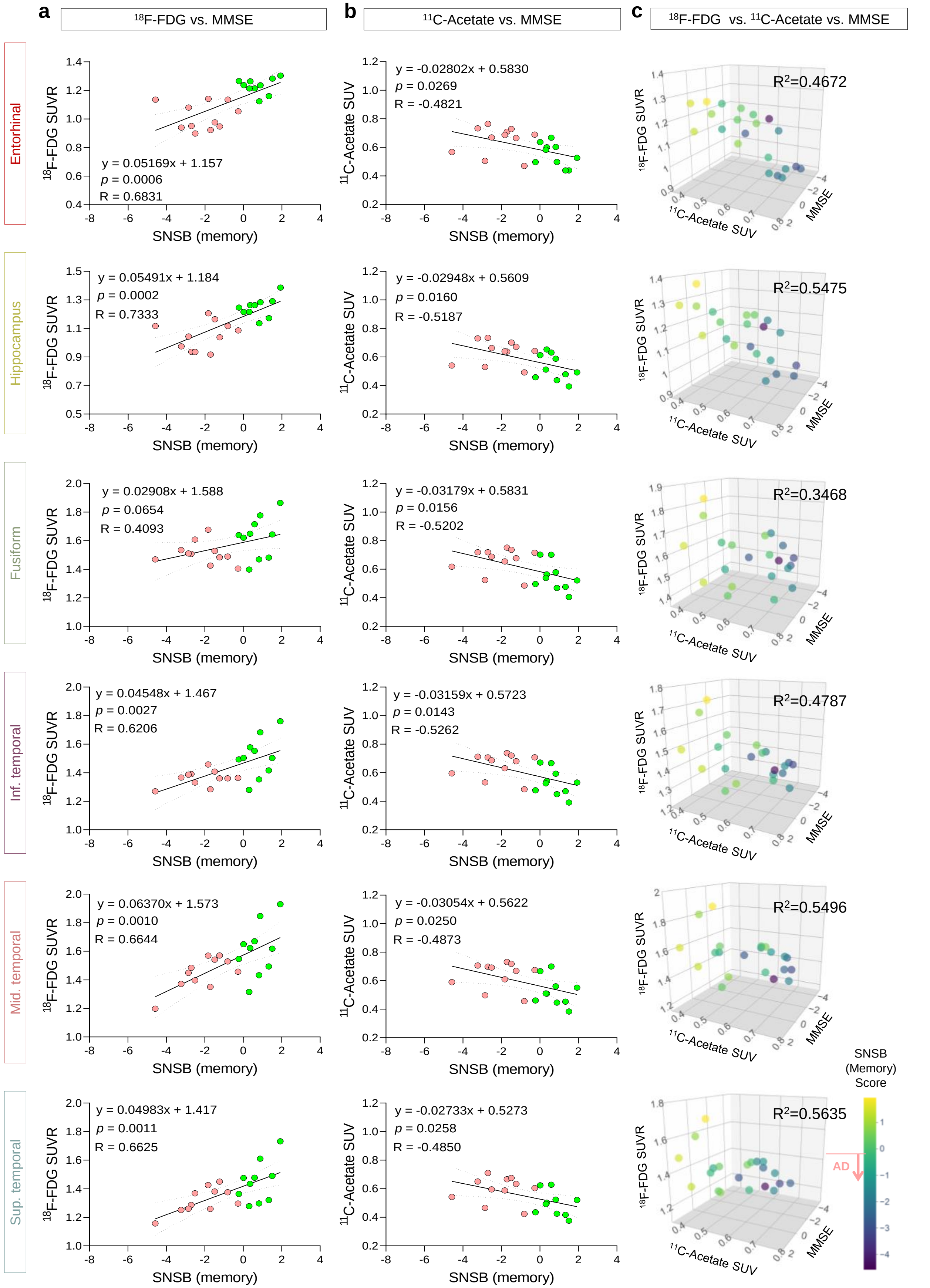

a
$^{18}$F-FDG vs. MMSE
b
$^{11}$C-Acetate vs. MMSE
c
$^{18}$F-FDG vs. $^{11}$C-Acetate vs. MMSE
Entorhinal
y = 0.05169x + 1.157
p = 0.0006
R = 0.6831
y = -0.02802x + 0.5830
p = 0.0269
R = -0.4821
$R^2$=0.4672
Hippocampus
y = 0.05491x + 1.184
p = 0.0002
R = 0.7333
y = -0.02948x + 0.5609
p = 0.0160
R = -0.5187
$R^2$=0.5475
Fusiform
y = 0.02908x + 1.588
p = 0.0654
R = 0.4093
y = -0.03179x + 0.5831
p = 0.0156
R = -0.5202
$R^2$=0.3468
Inf. temporal
y = 0.04548x + 1.467
p = 0.0027
R = 0.6206
y = -0.03159x + 0.5723
p = 0.0143
R = -0.5262
$R^2$=0.4787
Mid. temporal
y = 0.06370x + 1.573
p = 0.0010
R = 0.6644
y = -0.03054x + 0.5622
p = 0.0250
R = -0.4873
$R^2$=0.5496
Sup. temporal
y = 0.04983x + 1.417
p = 0.0011
R = 0.6625
y = -0.02733x + 0.5273
p = 0.0258
R = -0.4850
$R^2$=0.5635
$^{18}$F-FDG SUVR
$^{11}$C-Acetate SUV
SNSB (memory)
MMSE
SNSB (Memory) Score
AD

**Extended Data Figure 11. Correlation between $^{18}$F-FDG SUVR or $^{11}$C-Acetate SUV and SNSB (memory) score in various human brain region.**
**a, b**, In every region-of-interest, a significant correlation between $^{18}$F-FDG SUVR (**a**) or $^{11}$C-acetate SUV (**b**) and SNSB (memory) score was observed. **c,** The multiple linear re gression analysis showed that $^{18}$F-FDG SUVR and $^{11}$C-Acetate SUV has marked correl ations with SNSB (memory) scores.

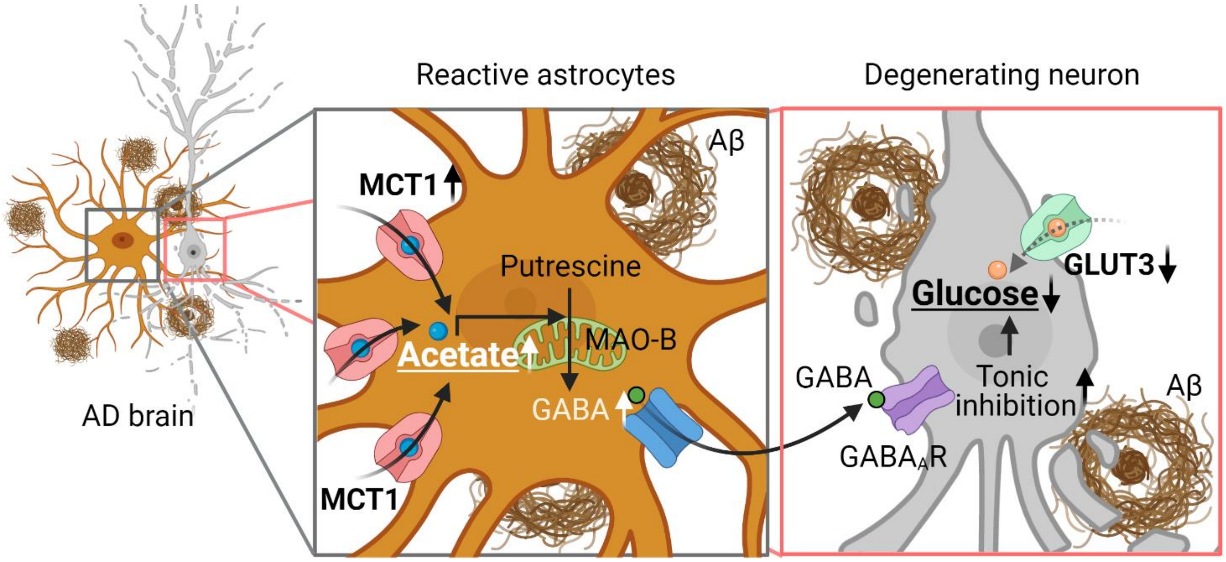


**Extended Data Figure 12. Schematic diagram of astrocytic acetate hypermetabolism and neuronal glucose hypometabolism.**
Excessive acetate metabolism of reactive astrocytes causes an aberrant synthesis of GABA, leading to reduced glucose metabolism of neighboring neurons in the AD brains.

**Extended Data Table 1. Information on postmortem brain tissue samples from normal subjects and AD patients.**

| Number | Case | Age | Sex | Braak stage |
|---|---|---|---|---|
| 1 | Normal | 87 | Female | I |
| 2 | Normal | 88 | Male | I |
| 3 | Normal | 87 | Female | II |
| 4 | Normal | 67 | Male | I |
| 5 | Normal | 82 | Male | I |
| 6 | Normal | 78 | Female | I |
| 7 | Normal | 70 | Male | I |
| 8 | Normal | 82 | Male | I |
| 9 | Normal | 79 | Female | I |
| 10 | Normal | 80 | Female | I |
| 1 | AD | 85 | Male | V |
| 2 | AD | 87 | Female | V |
| 3 | AD | 82 | Male | V |
| 4 | AD | 79 | Female | VI |
| 5 | AD | 70 | Male | VI |
| 6 | AD | 80 | Female | V |
| 7 | AD | 75 | Male | V |
| 8 | AD | 83 | Male | VI |
| 9 | AD | 79 | Female | VI |
| 10 | AD | 69 | Male | VI |

**Extended Data Table 2. Detailed information on human subjects for PET/CT imaging study**

| | Control | AD | ***p*** |
|---|---|---|---|
| n | 10 | 11 | |
| Demographics | | | |
| Age, yr | 69.00 ± 9.95 | 74.45 ± 6.47 | 0.169 |
| Sex, female (%) | 2 (20) | 7 (63.64) | 0.046 |
| Education, yr | 13.00 ± 3.79 | 8.18 ± 4.49 | 0.021 |
| Amyloid positivity, n | 0 | 7 | |
| Neuropsychological tests (SNSB) | | | |
| Attention function | 0.75 ± 0.97 | -0.85 ± 0.72 | <0.001 |
| Language function | 3.72 ± 1.96 | -0.88 ± 0.75 | <0.001 |
| Visuosptial function | 0.64 ± 0.69 | -1.10 ± 1.26 | 0.002 |
| Memory function | 0.75 ± 0.65 | -2.35 ± 1.03 | <0.001 |
| Frontal/executive function | 1.05 ± 0.96 | -2.03 ± 1.49 | <0.001 |
| K-MMSE score | 29.30 ± 0.64 | 19.45 ± 3.34 | <0.001 |
| CDR-SOB | 0.00 ± 0.00 | 0.73 ± 0.25 | <0.001 |
